# The metal binding site composition of the cation diffusion facilitator protein MamM cytoplasmic domain impacts its metal responsivity

**DOI:** 10.1101/2020.02.06.936617

**Authors:** Shiran Barber-Zucker, Anat Shahar, Sofiya Kolusheva, Raz Zarivach

**Affiliations:** Department of Life Sciences, Ben-Gurion University of the Negev, Beer Sheva 8410501, Israel; The National Institute for Biotechnology in the Negev, Ben-Gurion University of the Negev, Beer Sheva 8410501, Israel; Ilse Katz Institute for Nanoscale Science and Technology, Ben-Gurion University of the Negev, Beer Sheva 8410501, Israel

## Abstract

The cation diffusion facilitator (CDF) is a conserved family of divalent d-block metal cation transporters that extrude these cations selectively from the cytoplasm. CDF proteins are composed of two domains: the transmembrane domain, through which the cations are transported, and a regulatory cytoplasmic C-terminal domain (CTD). Metal binding to the CTD leads to its tighter conformation, and this sequentially promotes conformational change of the transmembrane domain which allows the actual transport of specific metal cations. It was recently shown that the magnetotactic bacterial CDF protein MamM CTD has a role in metal selectivity, as binding of different metal cations exhibits distinctive affinities and conformations. It is yet unclear whether the composition of the CTD binding sites can impact metal selectivity. Here we performed a mutational study of MamM CTD, where we exchanged the metal binding residues with different metal-binding amino acids. Using X-ray crystallography and Trp-fluorescence spectrometry, we studied the impact of the mutations on the CTD conformation in the presence of different metals. Our results reveal that the incorporation of such mutations alters the domain response to metals *in vitro*, as mutant forms of the CTD bind metals differently in terms of the composition of the binding sites and the CTD conformation.

**Coordinates:** MamM CTD structures have been deposited in the Protein Data Bank under the following accession codes: 6H5V, 6H5M, 6H5U, 6H8G, 6HAO, 6H88, 6H87, 6H8A, 6H89, 6H8D, 6H5K, 6H9Q, 6H84, 6H83, 6HA2, 6H8I, 6H9T, 6H81, 6HAN, 6H85, 6H9P, 6HHS.

## Introduction

Divalent d-block metal cations (DDMCs), such as Zn^2+^, Mn^2+^ and Fe^2+^, are crucial for numerous cell functions, hence their cellular accumulation should be effectively regulated^1,2^. The cation diffusion facilitator (CDF) proteins are a conserved family of DDMC transporters that extrude DDMCs from the cytoplasm through the cell membrane or into inner cellular compartments, usually by exploiting the proton motive force, and by so doing ensure the homeostasis of these cations at the cellular level^3^. Like other metalloproteins, each CDF protein can effectively bind and transport only specific metals^4–7^. In the case of CDF proteins, the metal selectivity mechanism should be highly sensitive, as these proteins control the overall concentrations of the DDMCs inside the cells and hence directly influence all metalloprotein function. CDF proteins form dimers and are typically composed of two domains^5,8–10^. The DDMCs are transported through the transmembrane domain via a conserved and well-defined monomeric metal binding site which also controls, to some extent, the metal selectivity^4,11,12^. The cytoplasmic C-terminal domain (CTD), which is frequently found in CDF proteins, forms a dimeric V-shape that enables the concerted movement of the full monomers, and serves as a regulatory domain^5,12–14^. In high DDMC concentrations, the DDMCs bind to specific metal binding sites in this domain, which causes a closure of the dynamic V-shaped apo form to a rigid and tighter conformation. This, in turn, facilitates the conformational change of the TMD from a cytoplasmic-facing conformation to an outer-membrane / inner-compartmental-facing conformation which allows the release of the DDMCs^5,12–15^. The CTD was also shown to be related with metal selectivity, which enables another level of regulation by this domain^16^. However, as the metal binding sites in this domain are not highly conserved, it is not clear whether the location and composition of the CTD binding sites can impact metal selectivity.

MamM is a CDF protein found in magnetotactic bacteria (MTB). MTB can sense the geomagnetic field and thereby navigate to their preferred habitats, usually oxi-anoxic zones in aquatic environments. The magneto-sensing property is enabled by the magnetosomes, sub-cellular organelles each composed of nanometer-sized iron-based magnetic particle (pure magnetite or gregite) enclosed in a protein-rich lipid membrane. In each bacterium, the magnetosomes are arranged in a chain-like fashion which creates the dipole moment needed for magnetic-reception^17–19^. MamM is a CDF iron transporter found in the magnetosome membrane that facilitates the accumulation of iron inside the magnetosomes, thus enabling the synthesis of the magnetic particles. It has been previously shown that the deletion of *mamM* gene, or even only of its CTD, abolishes magnetic particle formation^10^. Furthermore, mutations in its metal binding sites in both the TMD and CTD cause defective formation of the magnetic particles *in vivo*^10,14^. MamM CTD was well characterized *in vitro*: its crystal structure revealed that it has the characteristic fold of CDF proteins; the CTD structure was shown to be crucial for the overall protein function; DDMC binding to the CTD causes a conformational change from a dynamic-apo form to a rigid, more closed V-shaped structure; and it binds different DDMCs distinctively^14,16,20,21^. Comprehensive biophysical analysis of MamM CTD in the presence of Zn^2+^ and *in vivo* studies showed that MamM CTD dimer binds three ions by two binding sites: a central binding site composed of D249 and H285 from both monomers, and two symmetrical peripheral binding sites, each composed of H264 from one monomer and E289 from the second monomer^14,21^. The crystal structures of MamM CTD bound to Cu^2+^, Ni^2+^ and Cd^2+^ were previously determined, confirming the participation of the binding site residues in the chelation of the metals^16^. The Cu^2+^-bound structure shows tighter conformation compared to the apo form, with copper ions bound at the two sites by H285, H264 and E289 (through water molecule). The Cd^2+^-bound and Ni^2+^-bound structures showed no conformational changes compared to the apo form; however, Cd^2+^ is bound by D249 and H285 in the central site and Ni^2+^ by H264. Complementary isothermal titration calorimetry (ITC), pulsed electron-electron double resonance (PELDOR) spectroscopy and Trp-fluorescence spectrometry data showed that the varied metals indeed bind differently to the protein also in solution, in terms of number of binding sites, affinity and conformational changes, if at all, implying that the CTD has a direct role in metal selectivity^16^.

Here we investigated the impact of MamM CTD metal binding site composition on the CTD responsivity to different DDMCs, in terms of its binding ability and its related conformational changes. For that, we designed nine new mutated constructs of MamM CTD (residues 215-318), where we altered the binding site residues to different residues that tend to bind DDMCs. We created the D249H and H285D mutations to investigate whether a central binding site composed of four histidines or four aspartates can maintain the DDMCs binding abilities. Similarly, we created the H264E and E289H mutations for the investigation of the peripheral sites. To understand the importance of the location of the residues within the binding site for DDMCs binding, we also created two double mutations, D249H-H285D and H264E-E289H. Lastly, to examine the head group importance and influence on binding, we created the D249N, D249E and E289D mutations. All mutant constructs were studied in the presence of different DDMCs using X-ray crystallography and Trp-fluorescence spectrometry. The Trp-fluorescence spectrometry results reveal that each mutation has a different effect on the ability to bind the DDMCs in solution. The crystal structures of 22 mutants / mutant-DDMC pairs were detected and compared to the previously solved structures of MamM CTD with and without DDMCs, showing a range of possible conformations and binding sites that depend on both the nature of the substitution and the DDMC identity. Overall, our results show that the CDFs CTD metal binding sites might be altered to change the response of this domain to different DDMCs *in vitro*, and to impact the regulation of the whole protein. This study enables a rational design of MamM mutants so as to change its metal selectivity and to control its regulation, and this would constitute a first step for a future synthesis of magnetic particles with different chemical properties *in vivo*.

## Methods

### Site-directed mutagenesis and protein expression

*mamM* CTD gene from *Magnetospirillum gryphiswaldense* MSR-1 (UniProt Q6NE57 residues 215-318) was previously cloned into pET28a(+) vector (Novagen, Merck Biosciences, Germany)^22^; in this construct, pET28a-MamM-CTD-MSR1, the *mamM* gene was fused in-frame to express a six-His tag at the N-terminus of the protein followed by a thrombin proteolysis site. All MamM CTD mutations were applied to the pET28a-MamM-CTD-MSR1 vector using the QuickChange site-directed mutagenesis method (Stratagene, La Jolla, CA, USA). Primers containing single mutation sites (Hylabs, Rehovot, Israel) were designed and used for PCR amplifications. All MamM CTD forms were expressed similarly to those previously described for MamM CTD WT^22^.

### Protein purification

All MamM CTD constructs were purified as previously described for MamM CTD M250L and other mutants^20–22^. For all experiments, protein concentration was determined by measuring protein absorption at 280 nm.

### Crystallization and structure determination

Purified MamM CTD constructs at 20 mg mL^-1^ concentration in buffer containing 10 mM Tris pH=8.0, 150 mM NaCl, 5 mM β-mercaptoethanol and 3.375 mM metal solution (ZnCl_2_, CdCl_2_, MnCl_2_, NiCl_2_, FeCl_2_ or CuSO_4_), were subjected to crystallization trials using the vapor diffusion method at 293 K (0.3 μL protein with 0.3 μL reservoir solution for all protein-metal pairs). Crystals were harvested with or without treatment of cryo agent and flash-frozen in liquid nitrogen. Data collection was performed on a single-crystal at a temperature of 100 K. For all structures, data reduction was performed with XDS^23^ and data scaling with Aimless^24^ (after Aimless scaling, data of H264E Cd^2+^ form2, H264E-E289H, E289D Mn^2+^ form and E289D Cd^2+^ form were further processed with the STARANISO server (Global Phasing Ltd., Cambridge, UK). Phases were obtained by the molecular replacement method with the MamM CTD WT structure (PDB code: 3W5X^14^) as a template, using Phaser MR^25^. All structures were refined by Refmac5^26^, Phenix^27^ and/or PDB_REDO server^28^, while manual refinement was conducted using Coot version 0.8.9^29^. Aimless, Phaser MR and Refmac5 were used through the CCP4i package^30^. Rfree calculation used 5% of the data. All crystallization and cryo conditions, data collection details, data collection and refinement statistics and the used refinement software are given in Table S1-S3.

### Least-squares overlaps

All MamM CTD construct structures were overlapped and all figures were prepared using UCSF Chimera package, version 1.12^31^.

### Fluorescence spectrometry

Changes in tryptophan intrinsic fluorescence were recorded using Fluorolog®-3 (HORIBA Scientific, Edison, NJ, USA) equipped with quartz cell with 1 cm optical path length at ambient temperature. Samples of 1 mL MamM CTD proteins at 5 µM concentration in buffer containing 10 mM Tris pH=8.0 and 150 mM NaCl, were titrated using 2.5 mM metal solution in the same buffer to reach different concentrations (ZnCl_2_, CdCl_2_, NiCl_2_, MnCl_2_, CuSO_4_). Samples were measured at λex 297 nm, and the emission spectrum for each metal concentration was recorded at 310-450 nm. For each metal, the titration was replicated three times, and each spectrum was fitted to Extreme function by OriginPro (R-Square (COD) > 0.98) (OriginLab Corporation, Northampton, MA, USA). The maximum wavelength (wavelength at maximum intensity) and the intensity at that wavelength (aka maximum intensity) were averages for each metal concentration. Error is reported as the standard deviation. WT data is adapted from Barber-Zucker *et al.* 2020^16^. As Cd^2+^ and Zn^2+^ mainly cause concentration-dependent clear spectral shift and Ni^2+^ and Cu^2+^ concentration-dependent signal quenching, to simplify the analysis we discuss Cd^2+^ and Zn^2+^ impact on the spectral shift (hence their impact on the conformational change) and Ni^2+^and Cu^2+^ impact on the signal quenching (hence their ability to be bound closely to the W247 residue rather than their direct impact on the conformational change). The addition of Mn^2+^ did not result in signal quenching or spectral shift in any of the MamM CTD mutant constructs; hence, the Trp-fluorescence results with Mn^2+^ are not discussed in the *Results and Discussion* section.

## Results and Discussion

All nine designed mutants were well expressed and purified and were characterized in the presence of different DDMCs using X-ray crystallography and Trp-fluorescence spectrometry. To achieve better understanding on the conformational changes that occur due to DDMC binding in the different MamM CTD mutants, we attempted to crystallize MamM CTD mutants with diverse DDMCs under various conditions. We were able to successfully solve 22 structures of different MamM CTD mutants/ MamM CTD mutant-DDMC pairs (see Tables S1-S3 for more details). Among these structures, we detected the structures of all mutants; some in the presence of DDMCs, some in their apo forms and some in both states. We further characterized the differences in the binding abilities of all nine mutants and of the three binding-site deletion mutants: MamM CTD D249A-H285A (deletion of the central binding site), H264A-E289A (deletion of the peripheral binding site) and D249A-H264A (deletion of both sites) using Trp-fluorescence spectrometry, as previously described^14,16,21^. Here we titrated DDMCs (Cd^2+^, Mn^2+^, Ni^2+^, Cu^2+^ and Zn^2+^) into MamM CTD mutant solutions and measured the concentration-dependent changes in the tryptophan emission spectra (see *Methods* for analysis description). Below, we discuss the main differences observed between peripheral site mutants with different DDMCs, central site mutants with different DDMCs, or between different mutants with the same DDMC, while considering all related results.

### Impact of peripheral site mutants on DDMC binding

The peripheral site mutants in which we were able to determine their X-ray structure in the presence of different DDMCs are MamM CTD H264E and MamM CTD E289D, hence only these mutants will be analyzed and discussed in this section.

#### MamM CTD H264E with different metals

We solved the crystal structures of MamM CTD H264E with ZnCl_2_ in two different space groups, with CdCl_2_ in two different space groups and in the presence of MnCl_2_ in the crystallization condition - without, however, detecting Mn^2+^ in the electron density map (and therefore we relate to this structure as the apo form) (Figure 1A). The Cd^2+^-bound structures exhibit two conformations. The first structure [space group (SG) C222_1_] exhibits the same conformation as the wildtype (WT) apo and H264E apo but with two Cd^2+^ ions bound by each monomer’s D249 and H285 residues (Figure 1A, B). However, the occupancy of each ion in the electron density map is half, suggesting that only one ion can be bound by each monomer in either of the two binding sites. Another Cd^2+^ ion was detected in the periphery of each monomer, bound in a way that seems to have no influence on the protein conformation (Figure 1B). The second structure (SG P3_1_21) exhibits much tighter dimerization compared to that of the apo forms, with two symmetrical binding sites that involve H285 from one monomer and H236 and E289 from the second monomer. In none of the structures does the E264 residue participate in the Cd^2+^ binding. Both Zn^2+^-bound structures (SGs C121 and P3_1_21) exhibit the same conformation and binding sites which are shared with the second Cd^2+^-bound structure (Figure 1A, C), suggesting that this might be a preferred conformation for both cations.

**Figure 1:**
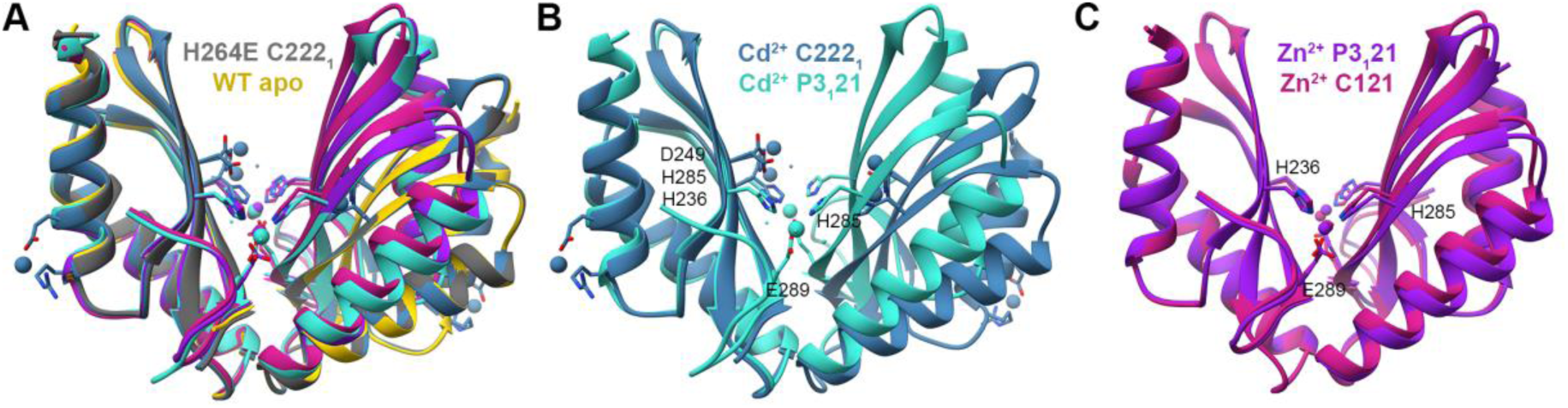
Crystal structures of MamM CTD H264E mutant with different metals. (A) Crystal structures of MamM CTD H264E with no metal bound (Mn^2+^ in crystallization condition, dim gray, pdb code: 6H5V), with Cd^2+^ (SG P3_1_21, turquoise, pdb code:6HAO; SG C222_1_, steel blue, pdb code: 6H8G) and with Zn^2+^ (SG P3_1_21, purple, pdb code:6H5M; SG C121, violet red, pdb code:6H5U) overlapped onto apo MamM CTD WT structure (SG C2221, gold, pdb code:3W5X^14^). (B) Crystal structures of MamM CTD H264E with Cd^2+^ at two different SGs (P3_1_21 turquoise, C222_1_ steel blue) show different binding sites and conformations of the mutant with Cd^2+^. (C) Crystal structures of MamM CTD H264E with Zn^2+^ at two different SGs (P3_1_21 purple, C121 violet red) show the same Zn^2+^ binding site which involves residues from both the central and peripheral binding sites, and the same closed conformation.

H264E-Cd^2+^ Trp-fluorescence scans show much smaller blue shift compared to the WT protein (and the smallest among all mutants; Figure 7B, upper panel), while H264E-Zn^2+^ Trp-fluorescence scans show some smaller blue shift compared to the WT protein (Figure 7A, upper panel), yet much larger compared to that of Cd^2+^. Nevertheless, previous PELDOR results of the WT CTD with Zn^2+^ and Cd^2+^ exhibit similar spin-labeling distances for both DDMCs, whereas the Cd^2+^-bound protein exhibits smaller blue-shift as compared to Zn^2+^, suggesting that Cd^2+^ causes smaller blue-shift although it undergoes similar closure in the presence of both DDMCs^16^.

**Figure 2:**
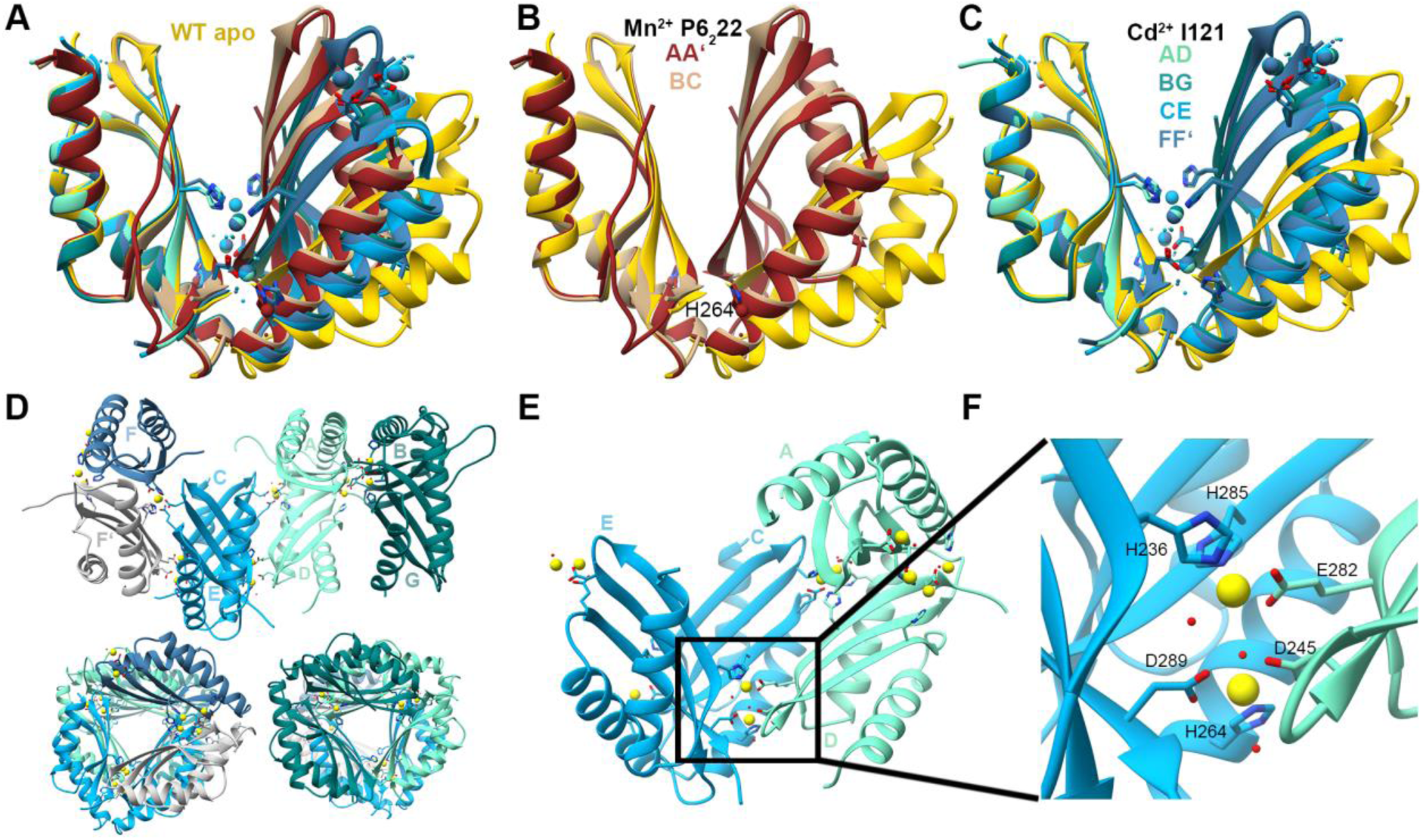
Crystal structures of MamM CTD E289D mutant with different metals. (A) Crystal structures of MamM CTD E289D with Cd^2+^ (SG I121, blue shades, pdb code: 6HHS) and Mn^2+^ (SG P6_2_22, brown shades, pdb code: 6H9P) overlapped onto apo MamM CTD WT structure (SG C222_1_, gold, pdb code: 3W5X^14^). As E289D-Cd^2+^ structure contains 7 monomers and E289D-Mn^2+^ structure contains 3 monomers in the asymmetric unit, all possible dimers are shown (four for the E289D-Cd^2+^ formed from the seven monomers and a monomer from an adjacent unit, and two for E289D-Mn^2+^, formed from the three monomers and a monomer from an adjacent unit). (B) Crystal structure of MamM CTD E289D with Mn^2+^ (chains A+A’ (tag for a monomer from an adjacent unit) in brown and chains B+C in tan) exhibits tighter conformation compared to the apo MamM CTD WT (gold). Mn^2+^ is bound by H264 and water molecule in each monomer of only the A+A’ dimer. (C) Crystal structure of MamM CTD E289D with Cd^2+^ (chains A+D in aquamarine, chains B+G in dark cyan, chains C+E in deep sky blue and chains F+F’ in steel blue) exhibits tighter conformation compared to the apo MamM CTD WT (gold), and different than that of E289D-Mn^2+^. Every Cd^2+^-pair is chelated by residues from three monomers (see Panels E+F); the upper Cd^2+^ pairs are shown for clarity to demonstrate how each monomer chelates the Cd^2+^ ions by three different sites. (D) Crystal packing of MamM CTD E289D with Cd^2+^ from three different angles. The asymmetric unit contains seven monomers that compose three biological dimers (chains A+D in aquamarine, chains B+G in dark cyan and chains C+E in deep sky blue). The seventh chain comprises an identical dimer with a monomer from an adjacent unit (chain F in steel blue and chain F’ from the adjacent unit in gray). (E) One dimer (chains C+E / chains A+D) and one monomer (chain D / chain C, respectively) are involved in the chelation of each Cd^2+^-pair. (F) Magnification of the Cd^2+^-pair binding site (Panel E, involving the chains C+D+E): H236 and D289 from chain E, H285 and H264 from chain C, D245 and E282 from chain D, and three water molecules, participate in the chelation of the Cd^2+^-pair.

**Figure 3:**
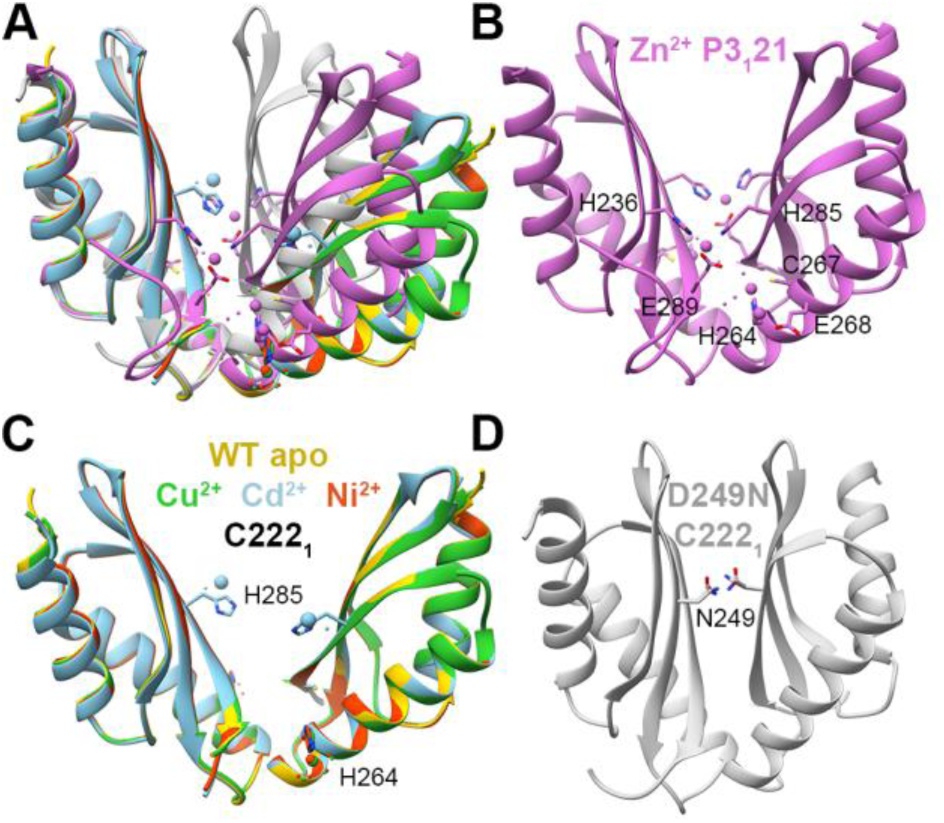
Crystal structures of MamM CTD D249N mutant with different metals. (A) Crystal structures of MamM CTD D249N with no metal bound (Mn^2+^ in crystallization condition, light gray, pdb code: 6H88), with Cd^2+^ (SG C222_1_, sky blue, pdb code: 6H8A), Cu^2+^ (SG C222_1_, lime green, pdb code: 6H89), Ni^2+^ (SG C222_1_, orange red, pdb code: 6H8D) and Zn^2+^ (SG P3_1_21, orchid, pdb code: 6H87), overlapped onto apo MamM CTD WT structure (SG C222_1_, gold, pdb code: 3W5X^14^). (B) Crystal structure of MamM CTD D249N with Zn^2+^ exhibits the same Zn^2+^ binding site as that of H264E-Zn^2+^ and the same conformation, with another two Zn^2+^ ions per monomer bound in two sites that involve H264. (C) Crystal structures of MamM CTD D249N with Cd^2+^ (sky blue), Cu^2+^ (lime green) and Ni^2+^ (orange red) show the same conformation as apo MamM CTD WT (gold). While Cu^2+^ and Ni^2+^ are chelated by H264 and water molecules, Cd^2+^ is bound by H285, water molecules, and H213 and E215 from a non-biological monomer from an adjacent unit (not shown). (D) Crystal structures of MamM CTD D249N with no metal bound (Mn^2+^ in crystallization condition) show very tight dimeric conformation that is stabilized by hydrogen bonds between the N249 residues from both monomers, which are replacing the charged and repulsive native-D249 in this mutant construct.

**Figure 4:**
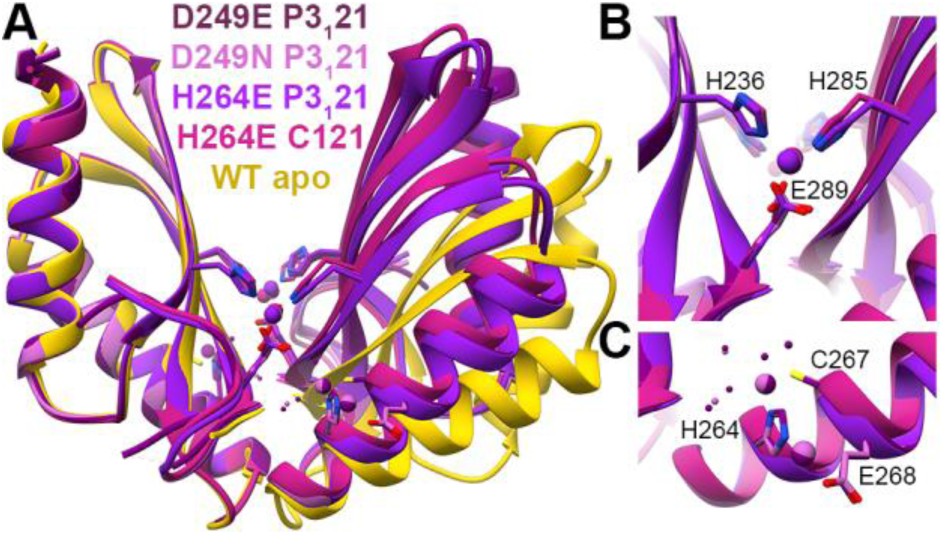
Crystal structures of different MamM CTD constructs bound to Zn^2+^. (A) Crystal structures of MamM CTD D249E with Zn^2+^ (SG P3_1_21, dark magenta, pdb code: 6H5K), MamM CTD D249N with Zn^2+^ (SG P3_1_21, orchid, pdb code: 6H87) and MamM CTD H264E with Zn^2+^ (SG P3_1_21, purple, pdb code: 6H5M; SG C121, violet red, pdb code: 6H5U) overlapped onto apo MamM CTD WT structure (SG C2221, gold, pdb code: 3W5X^14^). (B) The main Zn^2+^ binding site that is shared to all MamM CTD Zn^2+^-bound structures involves H236 and E289 from one monomer and H285 from the second monomer. (C) The peripheral binding site that is found only in MamM CTD D249E and D249N structures involves water molecules and residues only from one monomer (symmetric site): C267, E268 and H264. As can be evident in the H264E structures, the substitution of H264 to glutamate abolishes this site.

**Figure 5:**
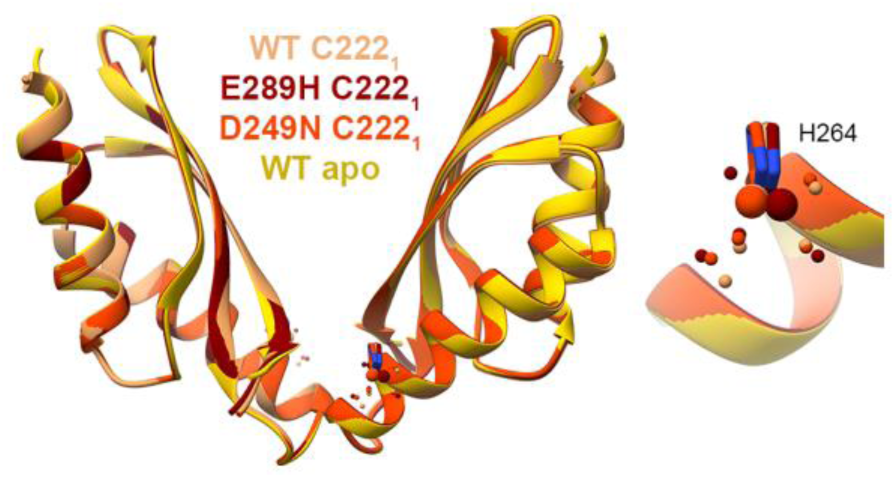
Crystal structures of different MamM CTD constructs bound to Ni^2+^: MamM CTD WT with Ni^2+^ (SG C222_1_, sandy brown, pdb code: 6GMV^16^), MamM CTD D249N with Ni^+^ (SG C222_1_, orange red, pdb code: 6H8D) and MamM CTD E289H with Ni^2+^ (SG C222_1_, dark red, pdb code: 6H81) overlapped onto apo MamM CTD WT structure (SG C2221, gold, pdb code: 3W5X^14^). All structures show the same conformation as the apo MamM CTD and bind Ni^2+^ by H264 and water molecules (right: enlarged binding site).

**Figure 6:**
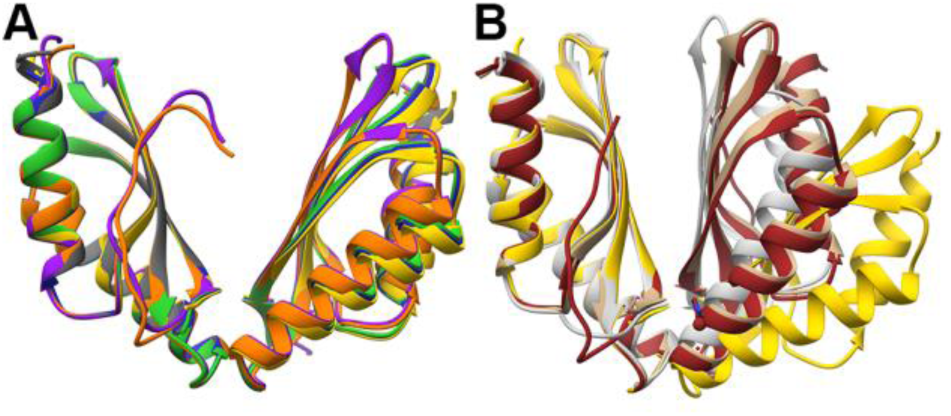
Crystal structures of different MamM CTD constructs that were crystallized in the presence of Mn^2+^. (A) Crystal structures of MamM CTD H285D (SG C222_1_, medium blue, pdb code: 6H8I), MamM CTD D249H-H285D (SG C222_1_, lime green, pdb code: 6HA2), MamM CTD H264E (SG C222_1_, dim gray, pdb code: 6H5V) and MamM CTD H264E-E289H (SG P2_1_2_1_2_1_, chains A+D in orange, chains B+C in purple, pdb code: 6HAN) overlapped onto apo MamM CTD WT structure (SG C2221, gold, pdb code: 3W5X^14^). All crystals were formed in the presence of Mn^2+^, however it could not be detected in the electron density map. Structures of all mutants have similar conformation to that of the WT apo. (B) Crystal structures of MamM CTD D249N (SG C222_1_, dim gray, pdb code: 6H88), and MamM CTD E289D (SG P6_2_22, chains A+A’ in brown, chains B+C in tan, pdb code: 6H9P) overlapped onto apo MamM CTD WT structure (SG C2221, gold, pdb code: 3W5X^14^). Both crystals were formed in the presence of Mn^2+^, however it could be detected in the electron density map of only one of the MamM CTD E289D dimers, chelated by only one residue, H264, and one water molecule. Both mutant structures exhibit tighter dimerization compared to the apo MamM CTD WT, yet the conformation is different between them.

**Figure 7:**
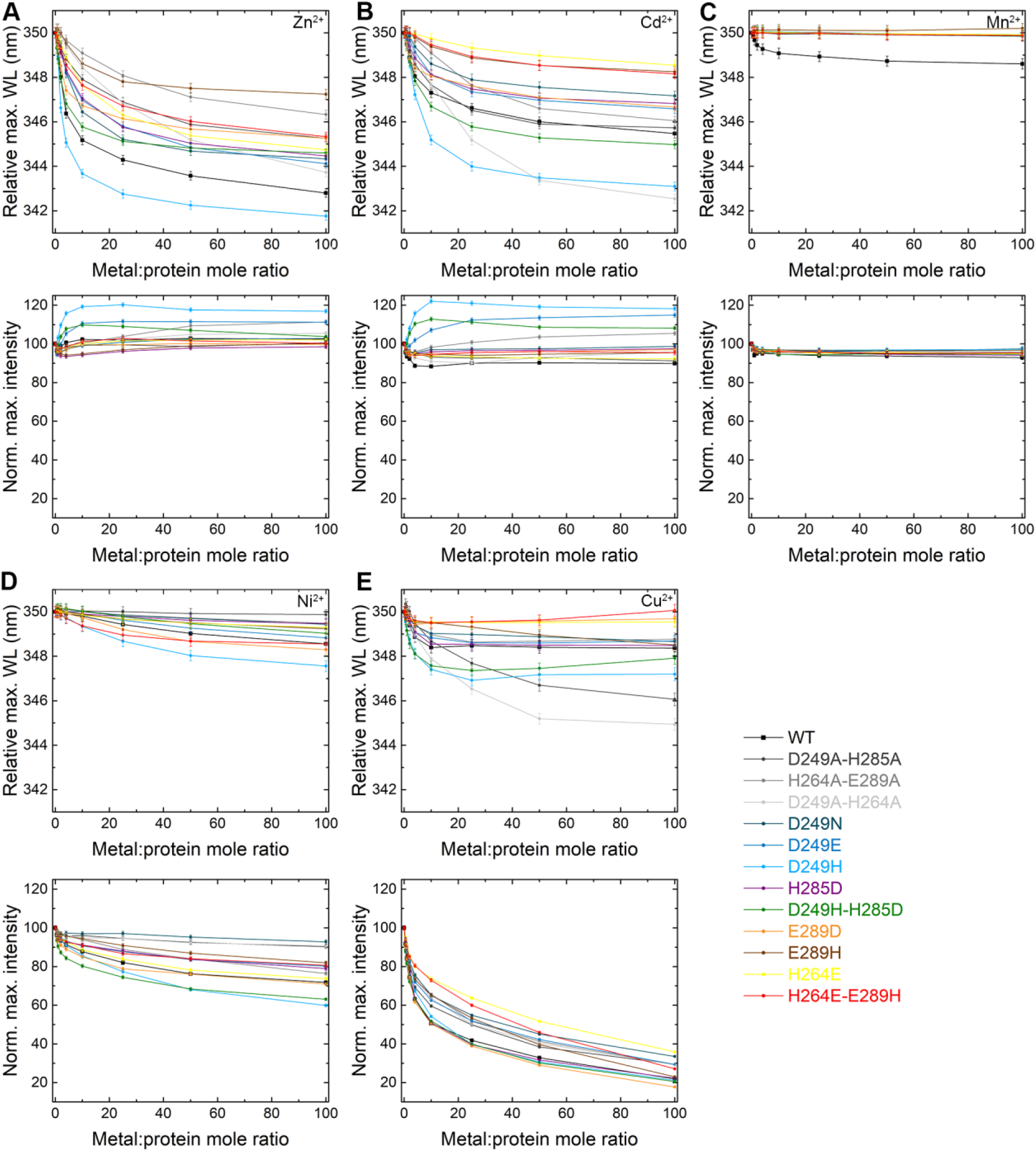
Fluorescence scans of MamM CTD constructs with varying metal concentrations. For A-E panels, the upper graph represents the maximum wavelength (normalized to 350 nm for easier comparison) as function of metal:protein ratio, while the lower panel represents the normalized fluorescence intensity compared to no metal as function of metal:protein ratio, of all MamM CTD constructs (WT, black, adapted from Barber-Zucker *et al.* 2020^16^; D249A-H285A, dark gray; H264A-E289A, gray; D249A-H264A, light gray; D249N, turquoise; D249E, dark blue; D249H, light blue; H285D, purple; D249H-H285D, green; E289D, orange; E289H, brown; H264E, yellow; H264E-E289H, red). MamM CTD proteins at 5 μM concentration were titrated using metal solutions: (A) Zn^2+^, (B) Cd^2+^, (C) Mn^2+^, (D) Ni^2+^, and (E) Cu^2+^, to reach different metal:protein ratios (intensity was normalized due to the change in MamM CTD concentration) and emission spectra were recorded. Samples were measured at an excitation of λex 297 nm and the emission spectrum for each metal concentration was recorded between 310-450 nm. For each protein, the data presented is the average of three independent measurements.

The calculated spin label distance of the apo WT protein is ∼ 7 Å longer than the evident WT Zn^2+^-bound distance^21^. This difference in the spin label distances was also shown for the apo and Cu^2+^-bound WT, suggesting that the WT binds Zn^2+^ and Cu^2+^ similarly and that the Cu^2+^-bound crystal structure represents also the conformation in solution^16^. However, the difference in distance between the same spin-labelled cysteine residues in the H264E-Zn^2+^ and the WT apo form is only ∼ 3 Å, which indicates a smaller conformational change as compared to that in the WT protein. Comparison of the Cu^2+^-bound WT structure to the H264E-Zn^2+^ bound structures reveals different orientation of the monomers, with the Cu^2+^-bound WT structure exhibiting much closer tryptophan residues with different environment as compared to the H264E-Zn^2+^ bound structures. Taking together all these crystallographic and fluorometric data, these results indicate that: (1) The addition of Cd^2+^ to H264E causes a minor conformational change in solution, if at all, and (2) Different conformational change occurs in H264E in the presence of Zn^2+^ compared to that in the WT, with the crystal structure probably representing also the conformation in solution.

#### MamM CTD E289D with different metals

We solved the crystal structures of MamM CTD E289D with CdCl_2_ and MnCl_2_ (Figure 2A). The E289D-Mn^2+^-bound structure contains three monomers in the asymmetric unit: two of them compose one biological dimer, and the third composes a biological assembly together with a monomer from an adjacent unit. Since a weak Mn^2+^ density was detected only in the latter dimer – bound to H264 and a water molecule – and as both dimers possess similar conformation (Figure 2B), it appears that the Mn^2+^ is bound non-specifically at that position. The conformation of this structure is much tighter than that of the WT apo form. In this structure, the D289 residue from one monomer forms a hydrogen bond with the second monomer H264, which stabilizes the closed conformation. In the WT protein the E289 residue could form this hydrogen bond, but as the glutamate residue is longer, the distance and angles between the other protomer’s residues would be different, which would lead to a smaller, less-stabilized network of bonds not sufficient for a steady closed conformation. This mutant is a great example of the dynamic conformation of the CTD in its unbound state, and of how the CTD conformation can be shifted by a seemingly minor mutation. The E289D-Cd^2+^-bound structure contains seven monomers in the asymmetric unit; six of them compose three biological dimers, and the seventh composes a biological assembly together with a monomer from an adjacent unit (Figure 2C-E). All four dimers exhibit the same Cd^2+^ binding sites and conformation, which is tighter than that of the WT apo form but less tight than that of the E289D-Mn^2+^-bound structure (Figure 2A, C). The dimer organization in the crystal (Figure 2D) reveals a unique Cd^2+^-pair binding site that involves residues from both dimer monomers and from a third monomer (that by itself form a dimer) (Figure 2E, F). The binding site involves residues from both central and peripheral binding sites; this binding site, and hence the conformation, could not exist in the presence of the longer E289 residue in the WT protein as it would deform the specific binding site geometry, showing again how this seemingly non-disruptive mutation can lead to different binding abilities. As this structure is not biological, it would not extend beyond this discussion.

The Trp-fluorescence maximum emission wavelength of E289D without the presence of any metals is shorter than that of the WT (Figure S2), although the residue is distant from W247 and therefore should not directly impact the fluorescence signal. This suggests that the initial conformation of the protein is tighter in this mutant form, as the E289D-Mn^2+^ structure also implies. The addition of Zn^2+^ and Cd^2+^ show some smaller blue-shift compared to the WT, while Ni^2+^ and Cu^2+^ show very similar trends in the signal quenching compared to the WT (Figure 7). However, as the initial maximum wavelength of E289D is shorter, it is not surprising to detect smaller shift in the wavelength; the E289D-Zn^2+^ and WT-Zn^2+^ maximum emission wavelengths at saturation are the same, hinting at a similar conformation, while that of E289D-Cd^2+^ is shorter than the WT-Cd^2+^ maximum emission wavelength at saturation, implying an even tighter conformation in this mutant form. For all other metals, as the signal quenching corresponds to that of the WT, one can expect similar conformations.

### Impact of central site mutants on DDMC binding

The peripheral site mutants in which we were able to determine their X-ray structure in the presence of different DDMCs are MamM CTD D249N, MamM CTD D249H and MamM CTD D249E. The crystal structures of MamM CTD D249E with CdCl_2_ and ZnCl_2_ and of MamM CTD D249H with CdCl_2_ and CuSO_4_ were solved, and their conformations and binding sites can be observed in Figure S3. As they have no unique conformations compared to the structures that are discussed in this section, they will be discussed only in the metal-based analysis below, therefore only the D249N mutant will be analyzed and discussed in this section.

#### MamM CTD D249N with different metals

We solved the crystal structures of MamM CTD D249N with ZnCl_2_, CdCl_2_, NiCl_2_ and CuSO_4_ and in the presence of MnCl_2_ in the crystallization condition, however without detecting Mn^2+^ in the electron density map (and therefore we relate to this structure as the apo form) (Figure 3A). The Zn^2+^-bound structure exhibits the same tighter conformation as the H264E-Zn^2+^ structure; nonetheless, it contains three symmetrical pairs of Zn^2+^ binding sites (Figure 3B). One of the binding sites is shared with the H264E-Zn^2+^ structure, while H264 residue binds the additional two Zn^2+^ ions in the periphery of each monomer, suggesting that the H264 residue has a role in attracting Zn^2+^ ions to the CTD. The Cd^2+^, Ni^2+^ and Cu^2+^ bound structures have the same conformation as the WT apo form (Figure 3C). While Ni^2+^ and Cu^2+^ are bound solely by H264 in each monomer (in addition to water molecules), one Cd^2+^ is chelated by H285 in each monomer, by water molecules and by H213 and E215 from a non-biological monomer from an adjacent unit. This implies that in contrast to the aspartate residue at position 249, asparagine cannot bind Cd^2+^. The last structure, that of the unbound D249N mutant, shows a very tight conformation compared to the WT apo and the D249N-Zn^2+^-bound structures (Figure 3A, D). This conformation is enabled by the D249N substitution that abolishes the original negative charge from the central binding site – which causes repulsion between the two monomers – and instead adds possible hydrogen bonds between the two asparagine residues that stabilize the closely-contacted monomers. Overall, the D249N crystal structures exhibit versatile conformation modes and binding sites that depend on the DDMC identity.

Trp-fluorescence results (Figure 7, S1 and S2), mainly of Ni^2 +^ and Cd^2+^ but also of Cu^2+^ and Zn^2+^ show smaller spectral shift (quenching/blue shift) when titrated to D249N, as compared to that of the WT. This means that Ni^2+^ and Cd^2+^ are not bound to the D249N mutant in a way that causes spectral shifts (in close proximity to the W247, and for Cd^2+^ also in a way that leads to conformational change). This is further supported by the crystal structures – where the Ni^2+^ is bound only in the periphery of the protein (by one residue which is distant from W247), and the Cd^2+^ is chelated only by H285 which is more distant than the D249 in the bound WT form^16^ – and where no conformational changes are shown. Similarly to H264E-Zn^2+^, that also exhibits the same crystallographic conformation, D249N-Zn^2+^ shows smaller blue-shift compared to WT and different binding site composition from WT^21^, indicating that it adopts the same conformation in solution as the crystal structure.

### DDMC binding to different mutants

For Zn^2+^, Cd^2+^, Ni^2+^ and Cu^2+^, we were able to determine the crystal structure of several mutants bound to each of these DDMCs. Here we focus on the analyses of the differences between the ability of different mutants to bind Zn^2+^, Ni^2+^ and Mn^2+^. Detailed analyses of Cd^2+^ and Cu^2+^ binding to the varied mutants is given in the *Supplementary Results and Discussion* section.

#### MamM CTD mutants binding to Zn^2+^

We solved the crystal structures of MamM CTD H264E (two different SGs), D249E and D249N with Zn^2+^. All structures exhibit the same fold and conformation, tighter than that of the WT apo (Figure 4A) and different from the Cu^2+^-bound WT structure, which is assumed to represent the Zn^2+^-bound structure as well. Examination of the Zn^2+^ binding sites reveals the importance of the H264 residue for Zn^2+^ binding. All three mutants have a shared symmetrical binding site, involving H285 from one monomer and H236 and E289 from the second monomer (Figure 4B). This binding site, that involves residues from both the central and peripheral binding sites, does not involve any of the mutated residues. The Zn^2+^-bound structure could not be obtained for the WT form. However, previous results suggest that the Zn^2+^ and Cu^2+^ lead to the same conformation and have the same binding sites^16,21^, suggesting that this site is not natural. The D249E and D249N mutants contain a second symmetrical binding site. In this site, the H264 residue chelates two Zn^2+^ ions together with C267 and E268 from the same monomer and water molecules (Figure 4C). This binding site seems not to influence the protein conformation, as it involves residues only from one monomer and as the H264E mutant shows the same fold. Nevertheless, the ability to bind four additional Zn^2+^ ions per dimer by H264 residue suggests that it might have a role in the attraction of ions to the CTD as an intermediate binding site, before the cations can be transported from this site to the second CTD binding site or to the TMD transport site. When comparing the Trp-fluorescence results of the WT and these mutants (Figure 7A, top panel), all three mutants show similar blue shift somewhat smaller than that of the WT, with the H264E showing the smallest shift of all. This indicates that all mutants have different and probably less tight conformation than that of the WT (as discussed above). Furthermore, it suggests that the inability to bind additional Zn^2+^ ions in the periphery of each monomer by H264 might lead to less stable closed conformation which results in smaller shift. Interestingly, D249H shows bigger shift than that of the WT, suggesting that although the 249 residues do not chelate the Zn^2+^ ions in the D249E/N mutants and Cu^2+^ in the WT Cu^2+^-bound structure, the additional histidine residue in D249H can help in the chelation of the Zn^2+^ to form a tighter conformation. This is supported by the fact that histidine residue tends to bind Zn^2+^ more than asparagine and glutamate^4^. In contrary, it might be that the bulky histidines lead to a different orientation of the monomers due to steric interference, and the bigger shift relates with a different swiveling movement of the monomers. All other mutants show smaller blue shift compared to the WT, implying the importance of every residue for proper metal binding.

#### MamM CTD mutant binding to Ni^2+^

For this study we solved the crystal structures of MamM CTD E289H and D249N with Ni^2+^, while we have previously solved the structure of the Ni^2+^-bound WT CTD^16^. All the Ni^2+^-bound structures exhibit the same conformation as the WT apo, with the Ni^2+^ ions bound only by each monomer’s H264 residue and water molecules (Figure 5). Previous PELDOR, Trp-fluorescence and ITC results of WT with Ni^2+^ suggested that Ni^2+^ binding is similar to that of Zn^2+^, where one ion is bound in the central site and two additional ions are bound to the peripheral sites, leading to a tight, rigid conformation as compared to the apo protein^16^. Because that the Ni^2+^-bound WT CTD crystal structure did not show the conformational changes expected from the studies in solution, we cannot conclude from the mutants’ crystal structures whether the mutations impact Ni^2+^-binding-dependent conformational changes. Contrary to the crystal structures, the Trp-fluorescence quenching patterns of the different mutant proteins with Ni^2+^ (Figure 7D, lower panel) show that each protein construct exhibits different binding abilities in solution. For example, the D249N, D249A-H264A and D249A-H285A show almost no quenching with the addition of Ni^2+^, suggesting that the D249 residue is important for Ni^2+^ binding near the W247 residues. D249H and D249H-H285D both show more quenching than that of the WT, and D249E shows less quenching, overall suggesting that it is not necessarily the aspartate in this location but that also histidine at the 249 position can bind Ni^2+^ in the central site. This is in agreement with the marked tendency of Ni^2+^ binding to histidine residues in crystal structures^4^. The E289H and E289H-H264E both show much less quenching compared to the WT, while E289D shows similar pattern to that of the WT. These Trp-fluorescence results demonstrate that the negative charge at position 289 is important for proper chelation in a distant central site, which also gives an indication of allostery between both sites as shown previously for Zn^2+^ binding^21^. H264E mutant, which was shown by the Zn^2+^, Cd^2+^ and Cu^2+^ Trp-fluorescence results to influence the binding and the correct conformational changes, shows the same pattern as the WT protein for Ni^2+^. This was a surprising result because the H264 residue is the only Ni^2+^-chelating residue in all structures, thereby strengthening evidence which points to this residue as an important factor for DDMC attraction. When considering all these results and the similarity between Zn^2+^ and Ni^2+^ binding in the WT CTD, we suggest that for Ni^2+^ as well the H264 position is not crucial for facilitation of the Ni^2+^ chelation in the central site – which would lead to the signal quenching – but only for the proper binding in the peripheral sites and the related conformational changes.

#### Unbound MamM CTD mutants and the special case of Mn^2+^

We solved the unbound structures of some MamM CTD mutants. All of them were crystallized in the presence of Mn^2+^; however, the Mn^2+^ could not be detected in their electron density map. MamM CTD H285D, D249H-H285D, H264E and H264E-E289H structures exhibit the same conformation as the WT apo form (Figure 6A). However, E289D and D249N exhibit much tighter conformation compared to the apo form, which differs from both structures (Figure 6B). As discussed above, one of the E289D dimers that could be built from the asymmetric unit monomers contained a weak manganese density, bound to H264 and water molecule; as the Mn^2+^ is bound solely at this position and could not be detected in the other dimer that adopts the same conformation, we assume that the Mn^2+^ is bound nonspecifically in this position and does not influence the conformational change. The reasons for the tighter conformation in both E289D and D249N apo forms were discussed above; both cases are great examples for how a single mutation can impact the dynamics of the apo forms, and potentially the regulation of the protein.

It was previously shown by Trp-fluorescence spectrometry, ITC and PELDOR that Mn^2+^ is the only DDMC out of six examined that MamM CTD WT protein could not bind^16^. We speculated that the mutants used in this study would show some binding abilities, as the binding site residues were confirmed and as Mn^2+^ is similar to the other DDMCs in its properties. Surprisingly, we could not detect Mn^2+^ in any of the crystal structures crystallized in its presence (except E289D where the Mn^2+^ is bound nonspecifically), and moreover, all mutants showed no Trp-fluorescence quenching (although paramagnetic) or blue shift in the presence of Mn^2+^ (Figure 7C). Mn^2+^ and Fe^2+^ are similar in their chemical properties. Interestingly, it was shown that uncultivated magnetotactic bacteria which are exposed to high concentrations of manganese can incorporate it into the iron-based magnetic particles^32^. Furthermore, another study demonstrated that Mn^2+^ has greater affinity to magnetite, both when it abiotically synthesized or in magnetotactic bacteria, as compared to other DDMCs^33^. To avoid overflow of manganese in the magnetosomes and a sequential incorporation of it into magnetite particles, we propose that MTB developed a sophisticated mechanism for selectivity against manganese. The selectivity that is observed by the CTD relates to more elements than to the binding sites residues, as we could not observe Mn^2+^ binding in any of our mutants, but we do not exclude the chance that other mutants would not revert to the observed selectivity.

## Conclusion

The DDMC binding site composition in MamM CTD impacts its ability to bind different DDMCs as well as impacting the bound-state conformation. The repertoire of bound conformations observed by the crystal structures is narrow, with all structures exhibiting the same or tighter conformation than the WT CTD. Furthermore, there are only a few specific residues that can participate in metal chelation by all mutant forms (as the DDMCs were not necessarily chelated by the binding site residues but also by other residues). These findings indicate that although we can alter the DDMC binding properties by mutating the binding site residues, in terms of conformation, identity of the binding sites and the number of metal bound to the CTD, the CTD conformational space and the possible binding sites are limited for only specific stable combinations. The CTD can control the metal selectivity of the overall protein by reacting distinctively to the different DDMCs; but only in response to some DDMCs can it adopt the exact tight conformation that will facilitate the sequential conformational change of the TMD that results in metal transport. The mutations can alter the conformational changes so that different monomer orientation or degree-of-closure of the CTD dimer are achieved in the presence of specific DDMCs so as to preserve, promote or block the actual transport. As these alterations putatively regulate the overall protein function, future studies will be able to use our results to design specific mutations in MamM *in vivo* to facilitate a controlled and specific DDMC binding to the CTD and, in combination with mutation in the transmembrane transport site, to insert DDMCs selectively into the magnetosome to form magnetic particles with varied magnetic properties.

## Supporting information

Tables S1-S3, Figure S1, Figure S2, Figure S3, Figure S4, Figure S5, Supplementary Results and Discussion

## Acknowledgments

SBZ and RZ are supported by the Israel Ministry of Science, Technology and Space, the Israel Science Foundation (grant no. 167/16), the European Molecular Biology Organization and CMST COST Action CM1306.

