## Supplementary material for "The metal binding site composition of the cation diffusion facilitator protein MamM cytoplasmic domain impacts its metal responsivity": Tables S1-S3, Figure S1, Figure S2, Figure S3, Figure S4, Figure S5, Supplementary Results and Discussion

In the main text we focused on the analyses of the differences between the ability of $Zn^{2+}$, $Ni^{2+}$ and $Mn^{2+}$ to be bound by different mutants (see main text *Results and Discussion*). Here we give detailed analyses of $Cu^{2+}$ and $Cd^{2+}$ binding to the varied mutants.

MamM CTD mutants binding to $Cu^{2+}$: For this study we solved the crystal structures of MamM CTD D249H and D249N with $Cu^{2+}$, while we previously solved the structure of the $Cu^{2+}$-bound WT CTD[1]. The WT bound structure shows tighter conformation compared to the WT apo form (Figure S4A), with $Cu^{2+}$ ions bound in the central binding site (one ion by H285 from both monomers) and peripheral binding sites (in each site, one ion by H264 from one monomer and a water molecule that bridges the E289 residue from the second monomer) (Figure S4B). Both D249H and D249N $Cu^{2+}$-bound structures exhibit the same conformation as the WT apo form, with the $Cu^{2+}$ only bound by the H264 residues and water molecules (Figure S4C). Although in the WT bound structure the D249 does not participate in the chelation of the $Cu^{2+}$ ions, it appears from the D249H and D249N $Cu^{2+}$-bound structures that this residue is important for the correct metal binding and conformational change. The bound $Cu^{2+}$ by H264 in the D249N/H structures again hint at the importance of this residue for the chelation of the $Cu^{2+}$ ions. Even if the H264 binding does not lead to conformational changes, it might act as an intermediate binding site. This is also supported by the Trp-fluorescence results of H264E, that show the least quenching of all mutants. In contrast to the crystal structures, the Trp-fluorescence results of the WT and the D249N/H mutants (Figure 7E, lower panel) show that D249H has similar quenching to the WT, while the D249N exhibits much less quenching. Hence, the asparagine in position 249 seems to inhibit the binding to the H285 residues, thereby causing less quenching, while the histidine does not. This suggests that in the case of D249H, the crystal structure does not accurately represent the state in solution. $Cu^{2+}$ was shown to be bound almost exclusively by histidine in crystal structures[2], and in the WT $Cu^{2+}$-bound structure it also appeared that the chelation by the histidine residues mostly contributes to the conformation stability. Hence, we propose that in contrast to the N249 residue in the D249N mutants, the H249 residue contributes to the stabilization of the $Cu^{2+}$ binding in the central site and to the closed conformation, although not observed in the crystal structure. The Trp-fluorescence results of D249H-H285D show also the same quenching pattern as in the WT and D249H, suggesting that indeed a histidine residue in the central site is important for proper chelation and conformational change.

MamM CTD mutants binding to $Cd^{2+}$: For this study we solved the crystal structures of MamM CTD D249H, D249E, D249N, H285D, H264E (two different SGs), H264E-E285H and E289D with $Cd^{2+}$; we previously solved the structure of the $Cd^{2+}$-bound WT CTD[1]. Only two $Cd^{2+}$-bound structures exhibit tighter conformation than the WT apo structure; one of the H264E structures and E289D (Figure S5A). Both structures exhibit the same conformation, but different binding site. While the H264E structure exhibits the same binding site as the H264E $Zn^{2+}$-bound structure (H285 from one monomer and H236 and E289 from the second monomer), the E289D exhibits a unique $Cd^{2+}$-pair binding site, as discussed in detail in the main text. All other MamM CTD constructs with $Cd^{2+}$, including the WT, adopt the same conformation as the WT apo, and the $Cd^{2+}$ is bound only in the central binding site by the residues in positions 249, 285 or both (Figure S5B). The exact $Cd^{2+}$ coordination depends on the substitution identity. In most mutants there is only one $Cd^{2+}$ ion bound to each monomer, while in others (WT and H264E) there are two ions that populate the binding site, but in half occupancy, suggesting that only one ion is effectively bound in them as well, by one of either binding sites. Furthermore, in some mutants the $Cd^{2+}$ is chelated by H213 and/or E215 from a non-biological monomer from an adjacent unit, which helps to form stable coordination of the $Cd^{2+}$ ions in the crystal form. Interestingly, in all structures where H285 residue exists it is involved in the chelation of the $Cd^{2+}$ ions, except in D249H where the H249 rather than H285 residue binds the ion. In the H285D mutant, the aspartate at position 249 binds the ion, overall showing that both the 249 and 285 positions and both histidine and aspartate residues can bind the $Cd^{2+}$ ions effectively. Previous PELDOR results show that the WT CTD binding to $Cd^{2+}$ lead to a tight rigid conformation, similar to that of the WT CTD with $Zn^{2+}$ and $Cu^{2+}$, and in contrast to the $Cd^{2+}$-bound crystal structure[1]. This indicates that for $Cd^{2+}$, the crystal structures do not necessarily represent well the structures in solution. In some mutants, $Cd^{2+}$ binding might not lead to conformational changes (as discussed below). In mutants that do no exhibit conformational changes in the $Cd^{2+}$-bound structures but do in solution, it is possible that $Cd^{2+}$ binding to each monomer prevents the tighter association of the monomers due to the large size of the $Cd^{2+}$ ions. Such an ion would not afford correct movement and stable tight conformation due to charge repulsion and steric interference. When considering the Trp-fluorescence results (Figure 7B, top panel), one can see that in solution the conformational changes occur, with each construct exhibiting different conformation. For example, H264E shows no blue-shift but one of its crystal forms showed tighter conformation, suggesting that H264E $Cd^{2+}$-bound structure, that exhibits no conformational change, is the conformation in solution (as discussed in the main text). E289D, whose crystal structure shows tighter conformation, shows a smaller blue shift compared to the WT. In this case, since: (1) The E289D-$Cd^{2+}$ and the H264E-$Zn^{2+}$ structures exhibit similar dimerization, (2) The addition of $Cd^{2+}$ to E289D and the addition of $Zn^{2+}$ to H264E result in smaller blue-shift as compared to their addition to the WT CTD, and (3) $Zn^{2+}$-bound and $Cd^{2+}$-bound conformations in solution are similar for the WT CTD (as evident by PELDOR[1]), we propose that the crystal structure of E289D-$Cd^{2+}$ also represents well the structure in solution. D249H shows the largest blue-shift, suggesting that the binding to position 249 exclusively (this is the only form where the ion was bound only at this position) impacts more on the conformational changes or on the Trp signal (as position 249 is closer to W247 than position 285).

**Supplementary Tables**

Table S1: Crystallization of MamM CTD constructs with different metals.

| Protein name | **H264E** | **H264E – $Zn^{2+}$ form1** | **H264E – $Zn^{2+}$ form2** | **H264E – $Cd^{2+}$ form1** | **H264E – $Cd^{2+}$ form2** |
|---|---|---|---|---|---|
| PDB code | 6H5V | 6H5M | 6H5U | 6H8G | 6HAO |
| Crystallization conditions | 0.2M $(NH_4)_2SO_4$, 0.1M BIS-TRIS pH=5.5, 25% PEG 3350 | 0.5M $Mg(HCO_2)_2$, 0.1M HEPES pH=7.5 | 0.2M $(NH_4)_2SO_4$, 0.1M BIS-TRIS pH=6.5, 25% PEG 3350 | 0.2M $(NH_4)_2SO_4$, 0.1M BIS-TRIS pH=5.5, 25% PEG 3350 | 0.2M $MgCl_2$, 0.1M BIS-TRIS pH=6.5, 25% PEG 3350 |
| Metal salt used (3.375 mM) | $MnCl_2$ | $ZnCl_2$ | $ZnCl_2$ | $CdCl_2$ | $CdCl_2$ |
| Cryo protectant | 50% PEG 3350 | 60% crystallizatio n condition, 15% MQ, 25% ethylene glycol | 50% PEG 3350 | - | 50% PEG 3350 |
| Data collection | ESRF – ID30B | ESRF – ID30B | ESRF – ID30B | ESRF – ID29 | BESSY II – BL 14.1 |
| Detector | Pilatus3 6M | Pilatus3 6M | Pilatus3 6M | Pilatus 6M | Pilatus 6M |

| Protein name | **D249N** | **D249N – $Zn^{2+}$ form** | **D249N – $Cd^{2+}$ form** | **D249N – $Cu^{2+}$ form** | **D249N – $Ni^{2+}$ form** |
|---|---|---|---|---|---|
| PDB code | 6H88 | 6H87 | 6H8A | 6H89 | 6H8D |
| Crystallization conditions | 0.1M HEPES pH=7.5, 25% PEG 3350 | 0.3M $Mg(HCO_2)_2$,0 .1M Tris pH=8.5 | 0.2M $Li_2SO_4$, 0.1M BIS-TRIS pH=5.5, 25% PEG 3350 | 0.2M $(NH_4)_2SO_4$, 0.1M BIS-TRIS pH=5.5, 25% PEG 3350 | 0.2M $Li_2SO_4$, 0.1M BIS-TRIS pH=5.5, 25% PEG 3350 |
| Metal salt used (3.375 mM) | $MnCl_2$ | $ZnCl_2$ | $CdCl_2$ | $CuSO_4$ | $NiCl_2$ |
| Cryo protectant | 50% PEG 3350 | 50% PEG 3350 | 50% PEG 3350 | - | - |
| Data collection | ESRF – ID29 | ESRF – ID29 | ESRF – ID29 | ESRF – ID29 | ESRF – ID29 |
| Detector | Pilatus 6M | Pilatus 6M | Pilatus 6M | Pilatus 6M | Pilatus 6M |

| Protein name | **D249E – $Zn^{2+}$ form** | **D249E – $Cd^{2+}$ form** | **D249H – $Cd^{2+}$ form** | **D249H – $Cu^{2+}$ form** | **D249H-H285D** |
| --- | --- | --- | --- | --- | --- |
| PDB code | 6H5K | 6H9Q | 6H84 | 6H83 | 6HA2 |
| Crystallization conditions | 0.3M $Mg(HCO_2)_2$,0.1M Tris pH=8.5 | 0.2M $Li_2SO_4$, 0.1M BIS-TRIS pH=5.5, 25% PEG 3350 | 0.2M $(NH_4)_2SO_4$, 0.1M BIS-TRIS pH=5.5, 25% PEG 3350 | 0.2M $(NH_4)_2SO_4$, 0.1M BIS-TRIS pH=5.5, 25% PEG 3350 | 0.1M NaOAc, pH=4.5, 25% PEG 3350 |
| Metal salt used (3.375 mM) | $ZnCl_2$ | $CdCl_2$ | $CdCl_2$ | $CuSO_4$ | $MnCl_2$ |
| Cryo protectant | 60% crystallization condition, 15% MQ, 25% ethylene glycol | - | - | 50% PEG 3350 | 50% PEG 3350 |
| Data collection | ESRF – ID30B | ESRF – ID29 | ESRF – ID23-1 | ESRF – ID23-1 | BESSY II – BL 14.1 |
| Detector | Pilatus3 6M | Pilatus 6M | Pilatus 6M | Pilatus 6M | Pilatus 6M |

| Protein name | **H285D** | **H285D – $Cd^{2+}$ form** | **E289H – $Ni^{2+}$ form** | **H264E-E289H** | **H264E-E289H – $Cd^{2+}$ form** |
| --- | --- | --- | --- | --- | --- |
| PDB code | 6H8I | 6H9T | 6H81 | 6HAN | 6H85 |
| Crystallization conditions | 0.2M $(NH_4)_2SO_4$, 0.1M BIS-TRIS pH=5.5, 25% PEG 3350 | 0.2M $Li_2SO_4$, 0.1M BIS-TRIS pH=5.5, 25% PEG 3350 | 0.2M $(NH_4)_2SO_4$, 0.1M BIS-TRIS pH=5.5, 25% PEG 3350 | 0.2M $(NH_4)_2SO_4$,0.1M HEPES pH=7.5, 25% PEG 3350 | 0.2M $(NH_4)_2SO_4$, 0.1M BIS-TRIS pH=5.5, 25% PEG 3350 |
| Metal salt used (3.375 mM) | $MnCl_2$ | $CdCl_2$ | $NiCl_2$ | $MnCl_2$ | $CdCl_2$ |
| Cryo protectant | 50% PEG 3350 | 50% PEG 3350 | 50% PEG 3350 | 50% PEG 3350 | 50% PEG 3350 |
| Data collection | ESRF – ID29 | ESRF – ID29 | ESRF – ID23-1 | BESSY II – BL 14.1 | DLS – I04 |
| Detector | Pilatus 6M | Pilatus 6M | Pilatus 6M | Pilatus 6M | Pilatus 6M-F |

| Protein name | **E289D – $Mn^{2+}$ form** | **E289D – $Cd^{2+}$ form** |
|---|---|---|
| PDB code | 6H9P | 6HHS |
| Crystallization conditions | 0.2M $Li_2SO_4$, 0.1M BIS-TRIS pH=5.5, 25% PEG 3350 | 0.1M $(NH_4)_2SO_4$, 0.1M BIS-TRIS pH=6.5, 0.1M NaCl |
| Metal salt used (3.375 mM) | $MnCl_2$ | $CdCl_2$ |
| Cryo protectant | 50% PEG 3350 | - |
| Data collection | ESRF – ID29 | BESSY II – BL 14.1 |
| Detector | Pilatus 6M | Pilatus 6M |

Table S2: Data collection and refinement statistics of MamM CTD constructs with different metals.

| Protein name | **H264E** | **H264E – $Zn^{2+}$ form1** | **H264E – $Zn^{2+}$ form2** | **H264E – $Cd^{2+}$ form1** | **H264E – $Cd^{2+}$ form2 [a]** |
|---|---|---|---|---|---|
| PDB code | 6H5V | 6H5M | 6H5U | 6H8G | 6HAO |
| Data collection | ESRF – ID30B | ESRF – ID30B | ESRF – ID30B | ESRF – ID29 | BESSY II – BL 14.1 |
| Space group | C 2 2 $2_1$ | P $3_1$ 2 1 | C 1 2 1 | C 2 2 $2_1$ | P $3_1$ 2 1 |
| ***Cell dimensions*** | | | | | |
| a, b, c (Å) | 37.18, 94.37, 53.09 | 53.27, 53.27, 61.42 | 88.05, 55.65, 63.77 | 37.00, 94.50, 53.21 | 53.66, 53.66, 58.07 |
| α, β, γ (°) | 90, 90, 90 | 90, 90, 120 | 90, 92.14, 90 | 90, 90, 90 | 90, 90, 120 |
| Resolution (Å) | 1.49-47.18 (1.49-1.52) | 1.60-46.14 (1.60-1.63) | 2.00-47.03 (2.00-2.05) | 1.35-47.25 (1.35-1.37) | 2.40-46.47 [b] (2.40-2.49) |
| Rsym or Rmerge | 0.065 (2.454) | 0.061 (1.898) | 0.083 (0.146) | 0.037 (1.443) | 0.169 (0.463) |
| I/σI | 13.9 (0.7) | 17.5 (1.3) | 23.2 (13.9) | 22.5 (1.3) | 11.9 (2.3) |
| $CC_{1/2}$ | 0.998 (0.356) | 0.999 (0.640) | 0.997 (0.993) | 0.999 (0.575) | 0.994 (0.910) |
| Completeness (%) | 99.6 (94.5) | 99.7 (92.3) | 99.8 (96.3) | 99.9 (99.1) | 98.6 (88.4) |
| Redundancy | 8.6 (7.5) | 12.7 (11.5) | 11.8 (12.0) | 7.1 (7.1) | 9.2 (4.7) |
| Wavelength (Å) | 0.97624 | 0.97624 | 0.97624 | 0.976247 | 0.9184 |
| No. unique reflections | 15616 | 13738 | 20715 | 20888 | 3988 |
| ***Refinement*** | | | | | |
| Resolution (Å) | 1.49-47.18 | 1.60-46.14 | 2.00-47.03 | 1.35-47.25 | 2.40-46.47 |
| Rwork/Rfree | 19.97/22.88 | 21.22/25.52 | 17.03/22.05 | 16.63/19.22 | 23.76/31.44 |
| *No. atoms* | | | | | |
| Protein | 644 | 688 | A-712, B-700, C-647, H-61 | 666 | 714 |
| Ligand/ion | 10 | 5 | 25 | 17 | 5 |
| Water | 49 | 25 | 159 | 53 | 11 |
| *B-factors* | | | | | |
| Protein | 35.11 | 43.25 | A-28.959, B-28.211, C-31.400, H-40.205 | 26.59 | 45.75 |
| Ligand/ion | $SO_4^{2-}$-35.49, βME-63.63, $Cl^-$-83.24 | $Zn^{2+}$-29.98, βME-55.92 | $SO_4^{2-}$-50.76, $Zn^{2+}$-16.44, βME-44.31 | $SO_4^{2-}$-38.18, $Cd^{2+}$-44.40, βME-51.75 | $Cd^{2+}$-32.92, βME-39.58 |
| Water | 40.42 | 45.85 | 33.44 | 38.82 | 33.00 |
| *RMSD* | | | | | |
| Bond lengths (Å) | 0.010 | 0.013 | 0.014 | 0.014 | 0.005 |
| Bond angles (°) | 1.437 | 1.617 | 1.467 | 1.624 | 0.900 |

| | | | | | |
|---|---|---|---|---|---|
| Ramachandran statistics [c] | P: 75 (97.40%), A: 1 (1.30%), O: 1 (1.30%) | P: 86 (98.85%), A: 1 (1.15%), O: 0 (0) | P: 251 (98.82%), A: 3 (1.18%), O: 0 (0) | P: 62 (98.41%), A: 1 (1.59%), O: 0 (0) | P: 83 (97.65%), A: 2 (2.235%), O: 0 (0%) |
| Missing residues | 293-318 | 211-212, 302-318 | A: 211, 302-318<br>B: 211, 303-318<br>C: 293-318<br>H: 211-299, 308-318 | 211-213, 293-318 | 302-318 |

| Protein name | **D249N** | **D249N – $Zn^{2+}$ form** | **D249N – $Cd^{2+}$ form** | **D249N – $Cu^{2+}$ form** | **D249N – $Ni^{2+}$ form** |
|---|---|---|---|---|---|
| PDB code | 6H88 | 6H87 | 6H8A | 6H89 | 6H8D |
| Data collection | ESRF – ID29 | ESRF – ID29 | ESRF – ID29 | ESRF – ID29 | ESRF – ID29 |
| Space group | C 2 2 $2_1$ | P $3_1$ 2 1 | C 2 2 $2_1$ | C 2 2 $2_1$ | C 2 2 $2_1$ |
| ***Cell dimensions*** | | | | | |
| a, b, c (Å) | 56.49, 62.83, 44.30 | 53.14, 53.14, 64.46 | 36.84, 94.66, 53.44 | 36.76, 94.63, 53.52 | 36.85, 94.33, 53.35 |
| α, β, γ (°) | 90, 90, 90 | 90, 90, 120 | 90, 90, 90 | 90, 90, 90 | 90, 90, 90 |
| Resolution (Å) | 1.50-42.01 (1.50-1.52) | 1.50-46.02 (1.50-1.52) | 1.80-47.33 (1.80-1.84 | 1.70-47.32 (1.70-1.73) | 1.62-47.16 (1.62-1.64) |
| Rsym or Rmerge | 0.038 (0.648) | 0.050 (0.753) | 0.145 (2.236) | 0.040 (1.064) | 0.033 (0.951) |
| I/σI | 22.2 (2.2) | 31.3 (2.9) | 12.3 (1.8) | 20.9 (1.4) | 23.2 (1.6) |
| $CC_{1/2}$ | 1.000 (0.814) | 1.000 (0.933) | 0.997 (0.666) | 0.999 (0.753) | 0.999 (0.828) |
| Completeness (%) | 99.2 (91.0) | 99.8 (95.3) | 100.0 (100.0) | 99.7 (95.0) | 99.7 (94.4) |
| Redundancy | 6.3 (5.6) | 12.6 (10.7) | 12.2 (12.8) | 6.4 (5.8) | 6.4 (6.0) |
| Wavelength (Å) | 0.975 | 0.975 | 0.975 | 0.975 | 0.975 |
| No. unique reflections | 12949 | 17417 | 9025 | 10686 | 12238 |
| ***Refinement*** | | | | | |
| Resolution (Å) | 1.50-42.01 | 1.50-46.02 | 1.80-47.33 | 1.70-47.32 | 1.62-47.16 |
| Rwork/Rfree | 18.21/22.66 | 18.93/22.71 | 19.24/23.67 | 19.59/24.58 | 17.30/21.36 |
| *No. atoms* | | | | | |
| Protein | 655 | 768 | 647 | 641 | 638 |
| Ligand/ion | 4 | 10 | 15 | 15 | 15 |
| Water | 74 | 81 | 55 | 51 | 63 |
| *B-factors* | | | | | |
| Protein | 21.21 | 28.31 | 30.25 | 33.21 | 32.02 |
| Ligand/ion | βME-32.16 | $Zn^{2+}$-21.14, βME-45.87, $CO_2$-53.56 | $SO_4^{2-}$-33.42, $Cd^{2+}$-35.43, βME-63.33 | $SO_4^{2-}$-34.79, $Cu^{2+}$-65.57, βME-50.32 | $SO_4^{2-}$-32.67, $Ni^{2+}$-67.79, βME-83.44 |
| Water | 32.26 | 33.98 | 37.75 | 41.19 | 41.96 |
| *RMSD* | | | | | |
| Bond lengths (Å) | 0.013 | 0.020 | 0.013 | 0.015 | 0.028 |
| Bond angles (°) | 1.658 | 2.034 | 1.592 | 1.609 | 2.558 |
| Ramachandran statistics [c] | P: 78 (97.50%), A: 2 (2.50%), O: 0 (0%) | P: 72 (98.63%), A: 1 (1.37%), O: 0 (0%) | P: 71 (98.61%), A: 1 (1.39%), O: 0 (0%) | P: 71 (98.61%), A: 1 (1.39%), O: 0 (0%) | P: 71 (98.61%), A: 1 (1.39%), O: 0 (0%) |
| Missing residues | 295-318 | 302-318 | 211-212, 293-318 | 211-212, 292-318 | 211-213, 293-318 |

| Protein name | **D249E – $Zn^{2+}$ form** | **D249E – $Cd^{2+}$ form** | **D249H – $Cd^{2+}$ form** | **D249H – $Cu^{2+}$ form** | **D249H-H285D** |
|---|---|---|---|---|---|
| PDB code | 6H5K | 6H9Q | 6H84 | 6H83 | 6HA2 |
| Data collection | ESRF – ID30B | ESRF – ID29 | ESRF – ID23-1 | ESRF – ID23-1 | BESSY II – BL 14.1 |
| Space group | P $3_1$ 2 1 | C 2 2 $2_1$ | C 2 2 $2_1$ | C 2 2 $2_1$ | C 2 2 $2_1$ |
| ***Cell dimensions*** | | | | | |
| a, b, c (Å) | 53.15, 53.15, 63.86 | 36.70, 94.52, 53.31 | 36.49, 94.52, 53.30 | 36.70, 95.07, 53.73 | 35.74, 94.30, 52.95 |
| α, β, γ (°) | 90, 90, 120 | 90, 90, 90 | 90, 90, 90 | 90, 90, 90 | 90, 90, 90 |
| Resolution (Å) | 1.54-46.03 (1.54-1.57) | 1.50-47.26 (1.50-1.52) | 2.04-47.26 (2.04-2.10) | 1.50-35.60 (1.50-1.52) | 1.50-47.15 (1.50-1.52) |
| Rsym or Rmerge | 0.115 (0.237) | 0.028 (0.963) | 0.128 (1.423) | 0.058 (1.735) | 0.046 (1.346) |
| I/σI | 20.8 (11.0) | 22.3 (1.3) | 13.9 (1.8) | 20.1 (1.4) | 18.6 (1.4) |
| $CC_{1/2}$ | 0.997 (0.990) | 1.000 (0.615) | 0.998 (0.603) | 0.999 (0.723) | 1.000 (0.582) |
| Completeness (%) | 99.9 (98.6) | 99.6 (94.1) | 99.7 (95.6) | 99.6 (91.6) | 99.7 (94.1) |
| Redundancy | 18.8 (18.5) | 4.3 (4.2) | 12.9 (10.7) | 13.0 (11.5) | 7.8 (7.3) |
| Wavelength (Å) | 0.97626 | 0.976247 | 0.977999 | 0.977999 | 0.9184 |
| No. unique reflections | 15927 | 15228 | 6119 | 15478 | 14775 |
| ***Refinement*** | | | | | |
| Resolution (Å) | 1.54-46.03 | 1.50-47.26 | 2.04-47.26 | 1.50-35.60 | 1.50-47.15 |
| Rwork/Rfree | 14.76/18.62 | 19.33/21.41 | 18.97/22.21 | 19.42/22.28 | 16.08/19.87 |
| *No. atoms* | | | | | |
| Protein | 715 | 646 | 639 | 653 | 640 |
| Ligand/ion | 7 | 15 | 15 | 16 | 4 |
| Water | 78 | 72 | 22 | 47 | 52 |
| *B-factors* | | | | | |
| Protein | 21.29 | 26.93 | 39.60 | 40.61 | 30.60 |
| Ligand/ion | $Zn^{2+}$-10.89, βME-36.12 | $SO_4^{2-}$-31.68, $Cd^{2+}$-24.04, βME-72.65 | $SO_4^{2-}$-46.19, $Cd^{2+}$-75.50, βME-78.26 | $SO_4^{2-}$-29.76, $Cu^{2+}$-61.52, βME-49.19, $Na^{+}$-57.50 | βME-42.57 |
| Water | 34.01 | 37.23 | 40.79 | 40.18 | 41.08 |
| *RMSD* | | | | | |
| Bond lengths (Å) | 0.011 | 0.017 | 0.016 | 0.013 | 0.015 |
| Bond angles (°) | 1.511 | 1.826 | 1.674 | 1.587 | 1.603 |
| Ramachandran statistics [c] | P: 84 (98.82%), A: 1 (1.18%), O: 0 (0) | P: 73 (98.65%), A: 1 (1.35%), O: 0 (0) | P: 78 (98.73%), A: 1 (1.27%), O: 0 (0%) | P: 71 (98.61%), A: 1 (1.39%), O: 0 (0%) | P: 71 (98.61%), A: 1 (1.39%), O: 0 (0%) |
| Missing residues | 293, 303-318 | 211-212, 293-318 | 293-318 | 211-212, 293-318 | 211-213, 293-318 |

| Protein name | **H285D** | **H285D – $Cd^{2+}$ form** | **E289H – $Ni^{2+}$ form** | **H264E-E289H** [a] | **H264E-E289H – $Cd^{2+}$ form** |
|---|---|---|---|---|---|
| PDB code | 6H8I | 6H9T | 6H81 | 6HAN | 6H85 |
| Data collection | ESRF – ID29 | ESRF – ID29 | ESRF – ID23-1 | BESSY II – BL 14.1 | DLS – I04 |
| Space group | C2 2 21 | C2 2 21 | C 2 2 $2_1$ | P $2_1$ $2_1$ $2_1$ | C 2 2 $2_1$ |
| ***Cell dimensions*** | | | | | |
| a, b, c (Å) | 36.44, 94.12, 53.07 | 36.37, 94.56, 53.30 | 37.06, 94.53, 53.54 | 65.59, 66.64, 113.29 | 36.23, 94.25, 53.10 |
| α, β, γ (°) | 90, 90, 90 | 90, 90, 90 | 90, 90, 90 | 90, 90, 90 | 90, 90, 90 |
| Resolution (Å) | 1.40-47.06 (1.40-1.42) | 1.70-47.28 (1.70-1.73) | 1.50-47.27 (1.50-1.53) | 2.60-46.75 [b] (2.60-2.71) | 2.00-53.10 (2.00-2.05) |
| Rsym or Rmerge | 0.036 (1.253) | 0.040 (0.829) | 0.049 (2.010) | 0.15 (1.191) | 0.051 (0.175) |
| I/σI | 21.4 (1.2) | 19.1 (1.6) | 22.6 (1.1) | 10.5 (1.4) | 29.1 (8.9) |
| $CC_{1/2}$ | 0.999 (0.639) | 1.000 (0.664) | 0.999 (0.611) | 0.997 (0.595) | 0.999 (0.993) |
| Completeness (%) | 99.9 (98.3) | 99.5 (98.5) | 99.5 (90.4) | 99.6 (97.3) | 99.6 (95.1) |
| Redundancy | 7.2 (6.6) | 4.2 (4.0) | 13.2 (10.8) | 8.4 (6.5) | 12.4 (10.2) |
| Wavelength (Å) | 0.976247 | 0.976247 | 0.977999 | 0.9184 | 0.9795 |
| No. unique reflections | 18410 | 10419 | 15422 | 15869 | 6470 |
| ***Refinement*** | | | | | |
| Resolution (Å) | 1.40-47.06 | 1.70-35.40 | 1.50-47.27 | 2.60-46.75 | 2.00-47.12 |
| Rwork/Rfree | 15.84/18.54 | 19.04/22.55 | 19.78/21.31 | 21.05/25.90 | 18.61/22.01 |
| *No. atoms* | | | | | |
| Protein | 665 | 659 | 632 | A-725, B-662, C-729, D-632 | 628 |
| Ligand/ion | 14 | 15 | 15 | 53 | 15 |
| Water | 59 | 54 | 39 | 56 | 28 |
| *B-factors* | | | | | |
| Protein | 27.01 | 31.72 | 37.22 | A-44.10, B-43.413, C-51.88, D-47.42 | 39.53 |
| Ligand/ion | $SO_4^{2-}$-30.98, βME-49.09 | $SO_4^{2-}$-35.85, $Cd^{2+}$-41.81, βME-71.72 | $SO_4^{2-}$-37.40, $Ni^{2+}$-84.22, βME-74.60 | $SO_4^{2-}$-44.88, βME-76.59, 1PE-37.31, $Cl^-$-48.61 | $SO_4^{2-}$-46.70, $Cd^{2+}$-46.92, βME-64.45 |
| Water | 37.98 | 39.60 | 43.60 | 35.28 | 41.40 |
| *RMSD* | | | | | |
| Bond lengths (Å) | 0.014 | 0.014 | 0.009 | 0.007 | 0.013 |

| | | | | | |
|---|---|---|---|---|---|
| Bond angles (°) | 1.658 | 1.573 | 1.458 | 1.116 | 1.606 |
| Ramachandran statistics [c] | P: 66 (98.51%), A: 1 (1.49%), O: 0 (0) | P: 71 (98.61%), A: 1 (1.39%), O: 0 (0) | P: 71 (98.61%), A: 1 (1.39%), O: 0 (0%) | P: 331 (96.50%), A: 8 (2.33%), O: 4 (1.17%) | P: 78 (98.73%), A: 1 (1.27%), O: 0 (0%) |
| Missing residues | 211-212, 293-318 | 211, 293-318 | 211-213, 293-318 | A: 211, 306-318, B: 295-318, C: 306-318, D: 293-318 | 211, 293-318 |

| Protein name | **E289D – $Mn^{2+}$ form [a]** | **E289D – $Cd^{2+}$ form [a]** |
|---|---|---|
| PDB code | 6H9P | 6HHS |
| Data collection | ESRF – ID29 | BESSY II – BL 14.1 |
| Space group | P $6_2$ 2 2 | I 1 2 1 |
| ***Cell dimensions*** | | |
| a, b, c (Å) | 78.26, 78.26, 175.50 | 83.60, 94.54, 107.49 |
| α, β, γ (°) | 90, 90, 120 | 90.00 91.76 90.00 |
| Resolution (Å) | 2.09-44.28 [b] (2.09-2.15) | 2.70-47.27 [b] (2.70-2.83) |
| Rsym or Rmerge | 0.081 (1.762) | 0.273 (1.117) |
| I/σI | 18.5 (1.2) | 5.7 (1.7) |
| $CC_{1/2}$ | 1.000 (0.837) | 0.984 (0.639) |
| Completeness (%) | 99.0 (87.6) | 99.3 (97.5) |
| Redundancy | 17.6 (13.8) | 6.8 (6.9) |
| Wavelength (Å) | 0.976247 | 0.9184 |
| No. unique reflections | 19491 | 22985 |
| ***Refinement*** | | |
| Resolution (Å) | 2.10 - 44.28 | 2.70-47.27 |
| Rwork/Rfree | 21.41/23.92 | 24.96/29.04 |
| *No. atoms* | | |
| Protein | A-723, B-617, C-630 | A-641, B-678, C-629, D-639, E-635, F-633, G-629, M-40 |
| Ligand/ion | 20 | 73 |
| Water | 86 | 113 |
| *B-factors* | | |
| Protein | A-39.93, B-43.63, C-54.34 | A-36.29, B-40.00, C-33.67, D-37.23, E-39.30, F-34.58, G-46.72, M-50.86 |

| | | |
|---|---|---|
| Ligand/ion | $SO_4^{2-}$-36.79, $Mn^{2+}$-76.74, βME-58.10 | $SO_4^{2-}$-35.78, $Cd^{2+}$-28.70, βME-53.39 |
| Water | 41.29 | 21.63 |
| *RMSD* | | |
| Bond lengths (Å) | 0.017 | 0.009 |
| Bond angles (°) | 1.904 | 1.346 |
| Ramachandran statistics [c] | P: 238 (99.58%), A: 1 (0.42%), O: 0 (0) | P: 559 (98.07%), A: 8 (1.40%), O: 3 (0.53%) |
| Missing residues | A- 303-318, B- 211-212, 292-318, C- 292-318 | A: 211-212, 296-318 B: 211-213, 302-318 C: 211-212, 294-318 D: 294-318 E: 211, 294-318 F: 211-212, 295-318 G: 211-212, 293-318 M: unknown residues |

Values in parentheses are for the highest resolution shell.

One crystal was used per dataset.

Data was collected at 100K.

[a] Collection statistics are given after the Aimless scaling[3]; data were further processed with the STARANISO server (http://staraniso.globalphasing.org/cgi-bin/staraniso.cgi; Global Phasing Ltd., Cambridge, UK).

[b] Resolution is given for best axis, as anisotropic data reduction in other axes resolution is worse.

[c] P- Preferred region, A- Allowed region, and O- outliers.

Table S3: Refinement software used for structure solution of MamM CTD constructs with different metals.

| Protein name | **PDB code** | **Refmac5**[4] | **Phenix**[5] | **PDB REDO**[6] |
|---|---|---|---|---|
| **H264E** | 6H5V | V | | V |
| **H264E – $Zn^{2+}$ form1** | 6H5M | V | | V |
| **H264E – $Zn^{2+}$ form2** | 6H5U | V | | V |
| **H264E – $Cd^{2+}$ form1** | 6H8G | V | | V |
| **H264E – $Cd^{2+}$ form2** | 6HAO | | V | V |
| **D249N** | 6H88 | | V | V |
| **D249N – $Zn^{2+}$ form** | 6H87 | V | V | V |
| **D249N – $Cd^{2+}$ form** | 6H8A | V | | V |
| **D249N – $Cu^{2+}$ form** | 6H89 | V | | V |
| **D249N – $Ni^{2+}$ form** | 6H8D | V | | |
| **D249E – $Zn^{2+}$ form** | 6H5K | V | | V |
| **D249E – $Cd^{2+}$ form** | 6H9Q | V | | V |
| **D249H – $Cd^{2+}$ form** | 6H84 | V | | V |
| **D249H – $Cu^{2+}$ form** | 6H83 | V | | V |
| **D249H-H285D** | 6HA2 | V | | V |
| **H285D** | 6H8I | V | | V |
| **H285D – $Cd^{2+}$ form** | 6H9T | V | | V |
| **E289H – $Ni^{2+}$ form** | 6H81 | V | | V |
| **H264E-E289H** | 6HAN | V | V | V |
| **H264E-E289H – $Cd^{2+}$ form** | 6H85 | V | | V |
| **E289D – $Mn^{2+}$ form** | 6H9P | V | | V |
| **E289D – $Cd^{2+}$ form** | 6HHS | V | V | V |

### Supplementary Figures

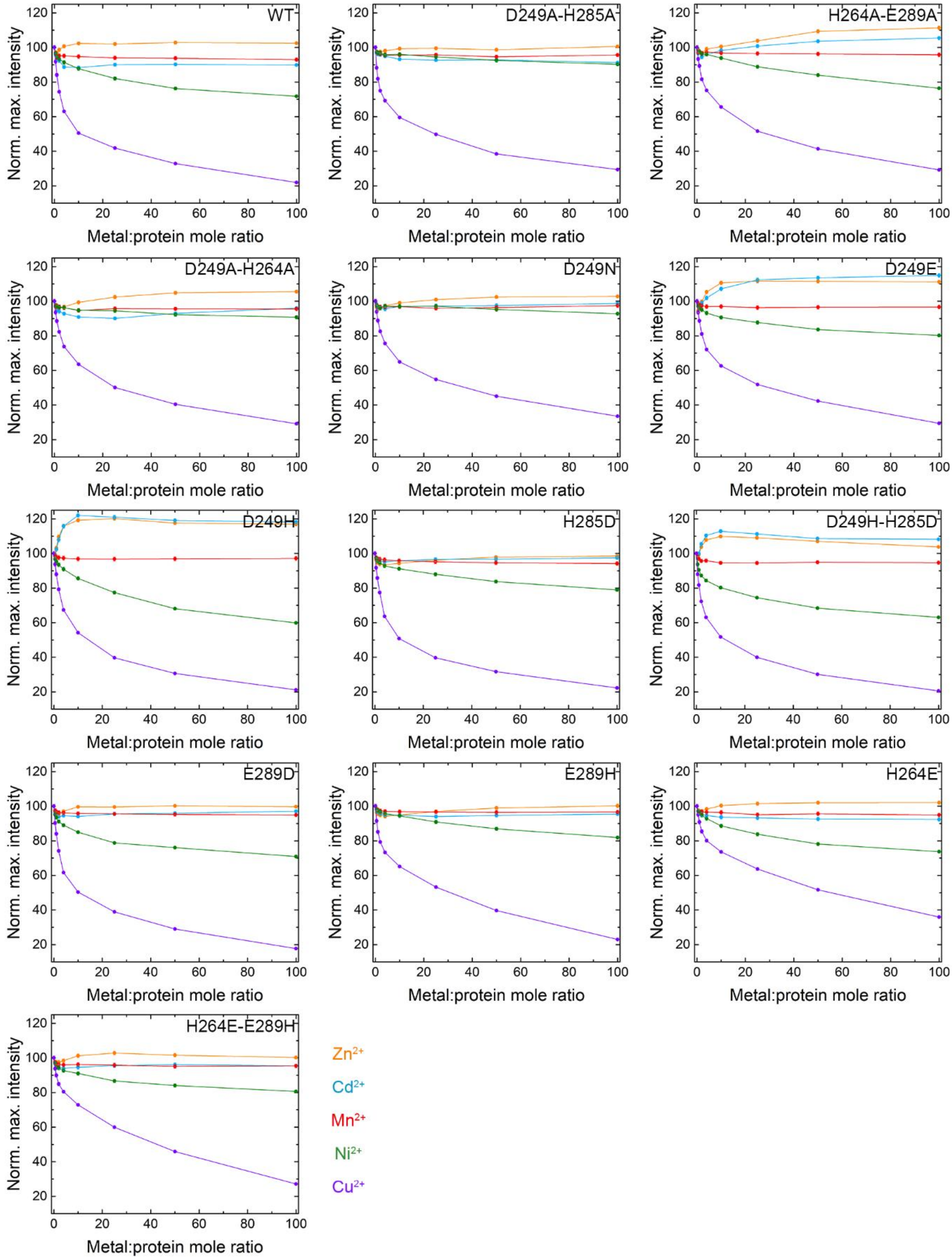


Figure S1: Normalized fluorescence intensity compared to no metal as function of metal:protein ratio of all MamM CTD constructs. Each graph represents one MamM CTD construct: WT (adapted from Barber-Zucker *et al.* 2020[1]), D249A-H285A, H264A-E289A, D249A-H264A, D249N, D249E, D249H,

H285D, D249H-H285D, E289D, E289H, H264E and H264E-E289H. MamM CTD proteins at 5 μM concentration were titrated using metal solutions ($ZnCl_2$, orange; $CdCl_2$, blue; $MnCl_2$, red; $NiCl_2$, green; $CuSO_4$, purple) to reach different metal:protein ratios (intensity was normalized due to the change in MamM CTD concentration) and emission spectra were recorded. Samples were measured at an excitation of λex 297 nm and the emission spectrum for each metal concentration was recorded between 310-450 nm. For each protein, the data presented is the average of three independent measurements.

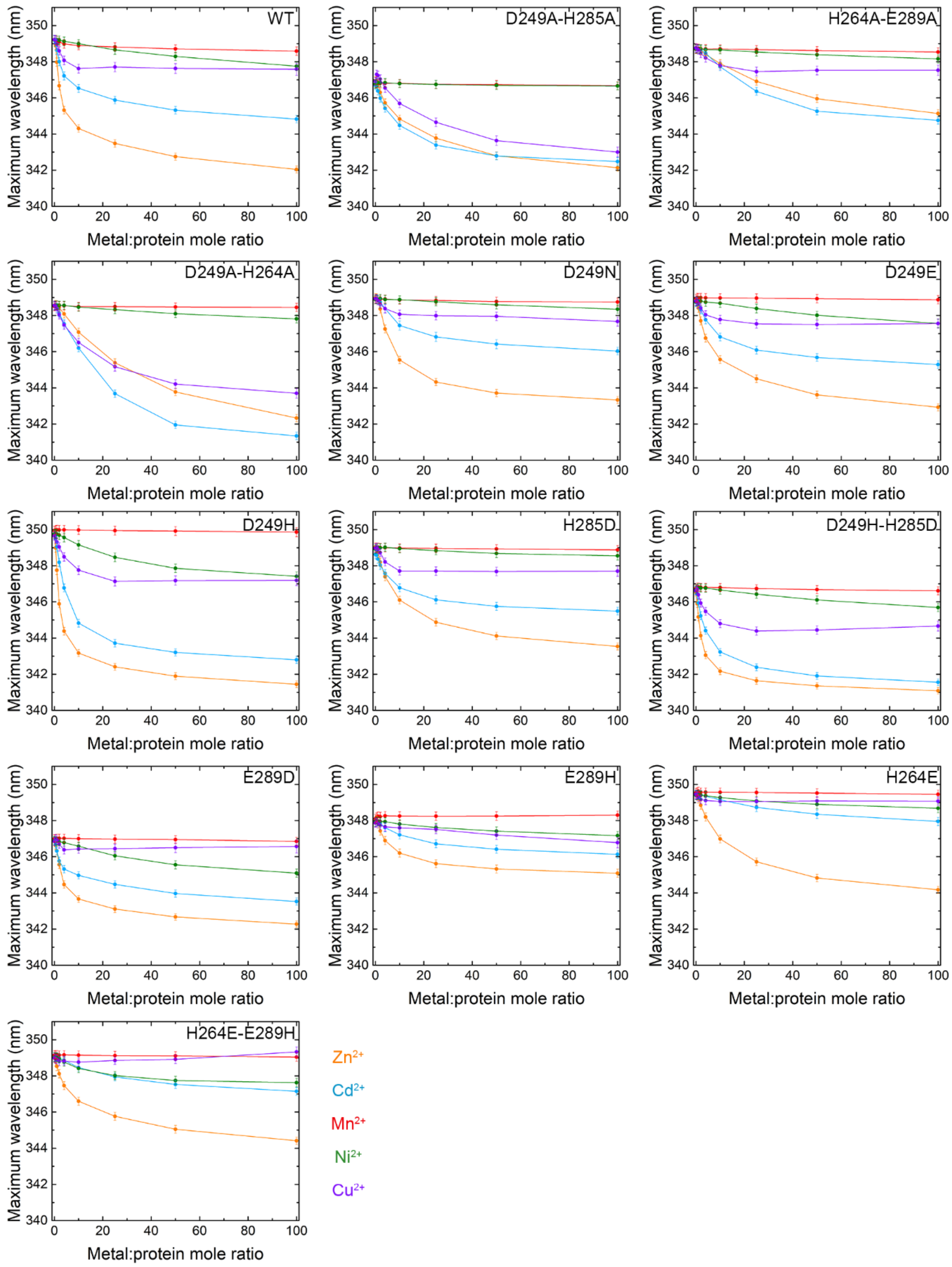


Figure S2: Maximum wavelength as function of metal:protein ratio of all MamM CTD constructs. Each graph represents one MamM CTD construct: WT (adapted from Barber-Zucker *et al.* 2020[1]), D249A-H285A, H264A-E289A, D249A-H264A, D249N, D249E, D249H, H285D, D249H-H285D, E289D, E289H, H264E and H264E-E289H. MamM CTD proteins at 5 µM concentration were titrated using metal solutions ($ZnCl_2$, orange; $CdCl_2$, blue; $MnCl_2$, red; $NiCl_2$, green; $CuSO_4$, purple) to reach different metal:protein ratios (intensity was normalized due to the change in MamM CTD concentration) and emission spectra were recorded. Samples were measured at an excitation of $\lambda$ex 297 nm and the emission spectrum for each metal concentration was recorded between 310-450 nm. For each protein, the data presented is the average of three independent measurements.

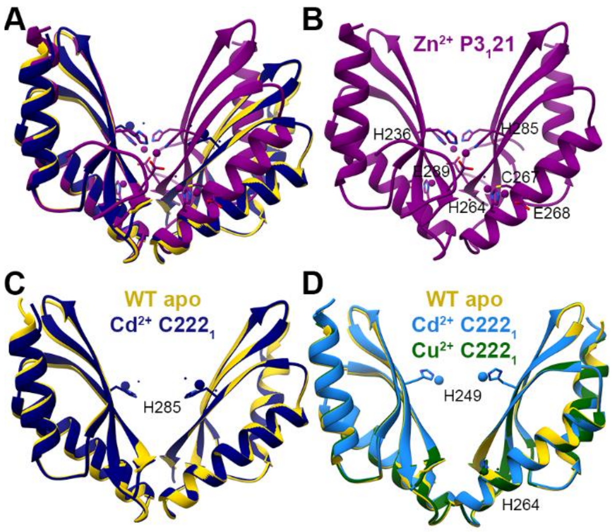


Figure S3: Crystal structures of MamM CTD D249E and D249H mutants with different metals. (A) Crystal structures of MamM CTD D249E with $Cd^{2+}$ (SG $C222_1$, navy blue, pdb code: 6H9Q) and $Zn^{2+}$ (SG $P3_121$, dark magenta, pdb code: 6H5K) overlapped onto apo MamM CTD WT structure (SG $C222_1$, gold, pdb code: 3W5X[7]). (B) Crystal structure of MamM CTD D249E with $Zn^{2+}$ exhibits the same $Zn^{2+}$ binding sites and conformation as that of D249N with $Zn^{2+}$. (C) Crystal structures of MamM CTD D249E with $Cd^{2+}$ (SG $C222_1$, navy blue) show the same conformation as apo MamM CTD WT (SG $C222_1$, gold). $Cd^{2+}$ is bound by H285, water molecules and H213 and E215 from a non-biological monomer from an adjacent unit (not shown). (D) Crystal structures of MamM CTD D249H with $Cd^{2+}$ (SG $C222_1$, dodger blue, pdb code: 6H84) and $Cu^{2+}$ (SG $C222_1$, dark green, pdb code: 6H83) overlapped onto apo MamM CTD WT structure (SG $C222_1$, gold, pdb code: 3W5X[7]). While $Cu^{2+}$ is chelated by H264 and water molecules, $Cd^{2+}$ is chelated by H249 and E215 from a non-biological monomer from an adjacent unit (not shown).

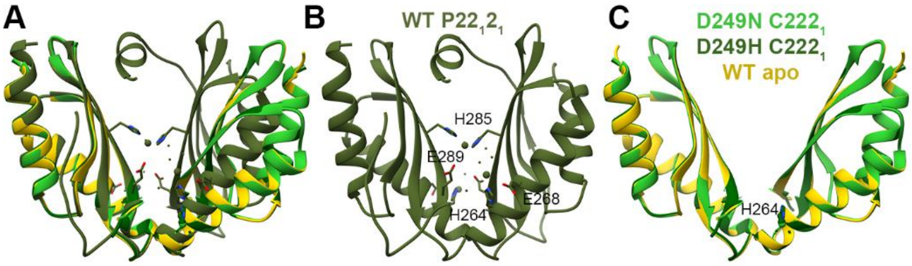


Figure S4: Crystal structures of different MamM CTD constructs bound to $Cu^{2+}$. (A) Crystal structures of MamM CTD WT with $Cu^{2+}$ (SG P22$_1$2$_1$, dark olive green, pdb code: 6GP6[1]), MamM CTD D249N with $Cu^{2+}$ (SG C222$_1$, lime green, pdb code: 6H89) and MamM CTD D249H with $Cu^{2+}$ (SG C222$_1$, dark green, pdb code: 6H83) overlapped onto apo MamM CTD WT structure (SG C2221, gold, pdb code: 3W5X[7]). (B) The crystal structure of MamM CTD WT with $Cu^{2+}$ exhibits tight conformation and contains one central $Cu^{2+}$ binding site and two symmetrical peripheral binding sites. (C) Crystal structures of MamM CTD D249N with $Cu^{2+}$ (SG C222$_1$, lime green) and MamM CTD D249H with $Cu^{2+}$ (SG C222$_1$, dark green) are bound only by H264 and water molecules and show the same conformation as apo MamM CTD WT (SG C222$_1$, gold).

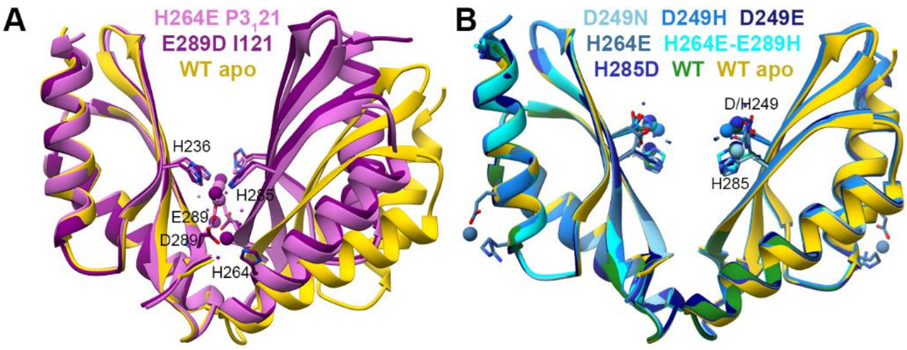


Figure S5: Crystal structures of different MamM CTD constructs bound to $Cd^{2+}$. (A) Crystal structures of MamM CTD H264E with $Cd^{2+}$ (SG P3$_1$21, orchid, pdb code: 6HAO) and MamM CTD E289D with $Cd^{2+}$ (SG I121, dark magenta, pdb code: 6HHS) overlapped onto apo MamM CTD WT structure (SG C2221, gold, pdb code: 3W5X[7]). Both structures exhibit similar conformation tighter than the WT apo, however the number of bound $Cd^{2+}$ ions per dimer and the binding sites differ between the two mutants (for E289D, two residues from a third monomer also participate in the chelation of each $Cd^{2+}$-pair, see Figure 2). (B) Crystal structures of MamM CTD with $Cd^{2+}$: H264E (steel blue, pdb code: 6H8G), H264E-E289H (cyan, pdb code: 6H85), D249N (sky blue, pdb code: 6H8A), D249E (navy blue, pdb code: 6H9Q), D249H (dodger blue, pdb code: 6H84), H285D (medium blue, pdb code: 6H9T) and WT (forest green, pdb code: 6GMT[1]) (all structures in SG C222$_1$) overlapped onto apo MamM CTD WT structure (SG C2221, gold, pdb code: 3W5X[7]). Although the number of bound $Cd^{2+}$ cations bound (one or two per monomer) and the exact residues that participate in the binding at each MamM CTD constructs differ, all structures show the same conformation and share similar binding properties (in most of the constructs, the $Cd^{2+}$ cations are also chelated by H213 and/or E215 from a non-biological monomer from an adjacent unit (not shown)).

**Supplementary Bibliographic References**


1 S. Barber-Zucker, J. Hall, A. Froes, S. Kolusheva, F. MacMillan and R. Zarivach, The cation diffusion facilitator protein MamM's cytoplasmic domain exhibits metal-type dependent binding modes and discriminates against $Mn^{2+}$. *bioRxiv*, 2020, preprint, DOI: 10.1101/2020.02.02.9.

2 S. Barber-Zucker, B. Shaanan and R. Zarivach, Transition metal binding selectivity in proteins and its correlation with the phylogenomic classification of the cation diffusion facilitator protein family. *Sci. Rep.*, 2017, **7**, 16381.

3 P. R. Evans and G. N. Murshudov, How good are my data and what is the resolution? *Acta Crystallogr. Sect. D Struct. Biol.*, 2013, **69**, 1204–1214.

4 G. N. Murshudov, P. Skubák, A. A. Lebedev, N. S. Pannu, R. A. Steiner, R. A. Nicholls, M. D. Winn, F. Long and A. A. Vagin, REFMAC5 for the refinement of macromolecular crystal structures. *Acta Crystallogr. Sect. D Struct. Biol.*, 2011, **67**, 355–367.

5 P. D. Adams, P. V Afonine, G. Bunkóczi, V. B. Chen, I. W. Davis, N. Echols, J. J. Headd, L.-W. Hung, G. J. Kapral, R. W. Grosse-Kunstleve, A. J. McCoy, N. W. Moriarty, R. Oeffner, R. J. Read, D. C. Richardson, J. S. Richardson, T. C. Terwilliger and P. H. Zwart, PHENIX : a comprehensive Python-based system for macromolecular structure solution. *Acta Crystallogr. Sect. D Struct. Biol.*, 2010, **66**, 213–221.

6 R. P. Joosten, F. Long, G. N. Murshudov and A. Perrakis, The PDB_REDO server for macromolecular structure model optimization. *IUCrJ*, 2014, **1**, 213–220.

7 N. Zeytuni, R. Uebe, M. Maes, G. Davidov, M. Baram, O. Raschdorf, M. Nadav-Tsubery, S. Kolusheva, R. Bitton, G. Goobes, A. Friedler, Y. Miller, D. Schüler and R. Zarivach, Cation diffusion facilitators transport initiation and regulation is mediated by cation induced conformational changes of the cytoplasmic domain. *PLoS One*, 2014, **9**, e92141.
